# Bayesian causal inference unifies perceptual and neuronal processing of center-surround motion in area MT

**DOI:** 10.1101/2025.09.17.676722

**Authors:** Gabor Lengyel, Sabyasachi Shivkumar, Gregory C. DeAngelis, Ralf M. Haefner

## Abstract

Center–surround interactions are a hallmark of visual processing and are especially prominent in area MT, where surround motion can either suppress or facilitate neuronal responses depending on context. However, existing mechanistic descriptions, including divisive normalization, do not explain the full diversity of these effects or their relationship to motion perception. Here, we show that both perceptual and neuronal center–surround phenomena can be understood as consequences of Bayesian causal inference over reference frames. Building on a normative model of motion perception, we derived predictions for the mean responses and variability of single MT neurons across the full four-dimensional space of center and surround directions and speeds. The model generates structured patterns of suppression, facilitation, and coordinate-frame selectivity that qualitatively match the diversity of center–surround effects reported in primate MT. Our results provide a unified computational account linking motion integration and segmentation in perception with contextual response modulation in MT, and yield testable predictions for how the visual system infers and represents reference frames.

## 1 Introduction

Center-surround processing is a canonical computation in the brain, shaping vision from its earliest stages (Allman et al., 1985b; Angelucci et al., 2017). Starting in the retina, where it enhances contrast sensitivity and facilitates the detection of boundaries (Baden et al., 2020; Euler et al., 2014; Gaynes et al., 2022; Kuffer, 1953; Turner et al., 2018), this principle of surround-dependent contextual modulation is repeated throughout the visual hierarchy. In the ventral stream, it refines feature selectivity in V1 (Angelucci et al., 2017; Nurminen & Angelucci, 2014) and shapes object representations in the inferotemporal cortex (Angelucci et al., 2017; Carandini & Heeger, 2012; Zoccolan et al., 2005). This principle is also fundamental to the dorsal stream’s processing of motion (Born, 2000; Born & Bradley, 2005; Britten & Heuer, 1999; DeAngelis & Uka, 2003; Pack et al., 2004; Rust et al., 2006). In the middle temporal area (MT), for instance, a moving surround can powerfully suppress a neuron’s response or, under different conditions, facilitate it (Allman et al., 1985a; Born, 2000; Born & Bradley, 2005; Born & Tootell, 1992; Bradley & Andersen, 1998; DeAngelis & Uka, 2003; Huang et al., 2007; Inaba et al., 2011, 2007; Newsome et al., 1988; Pack et al., 2004, 2005; Raiguel et al., 1995; Tanaka et al., 1986; Tzvetanov & Womelsdorf, 2008; Xiao et al., 1997; Zemel et al., 1998). Perceptually, this same stimulus can cause a central motion to be integrated with the surround into a unified whole object motion, or to be segmented from the surround in stark contrast (Bill et al., 2022, 2020; Braddick, 1993; Dakin & Mareschal, 2000; Gershman et al., 2016; Penaloza et al., 2024; Shivkumar et al., 2025; Tadin et al., 2003, 2019; Zarei Eskikand et al., 2020).

Despite their ubiquity, a unifying computational model explaining these diverse center-surround interactions on both at the perceptual and neural levels has remained elusive. At the neural level, we have a rich, process-level description of what happens, but no normative theory for why it happens and what computational principles can produce such profoundly different and context-dependent outcomes. For instance, decades of neurophysiological research have meticulously catalogued the variety of center-surround interactions in area MT, from the direction-selective antagonistic suppression first described by Allman et al. (1985a) and Tanaka et al. (1986), to the reinforcing facilitation and more complex patterns later categorized by Born (2000), Born and Bradley (2005), and Born and Tootell (1992). Remarkably, although some of these findings date back four decades, this diversity of effects, including both suppression and facilitation, still lacks a unified explanation. Existing models of the V1-MT pathway, often based on linear-nonlinear integration or divisive normalization, can account for a subset of suppressive phenomena but struggle to explain the full range of effects (Adelson & Bergen, 1985; Albright, 1984; Barth & Watson, 2000; Movshon et al., 1985; Qian et al., 1994a; Rust et al., 2006; Simoncelli & Heeger, 1998; Wilson et al., 1992; Wilson & Kim, 1994; Zarei Eskikand et al., 2020). Conversely, influential models of perception successfully describe phenomena like motion integration and segmentation but have not been linked to the rich diversity of single-neuron center-surround response properties (Bill et al., 2022, 2020; Braddick, 1993; Dakin & Mareschal, 2000; Gershman et al., 2016; Penaloza et al., 2024; Shivkumar et al., 2025; Tadin et al., 2003, 2019; Zarei Eskikand et al., 2020).

Recent normative models of motion perception suggest that causal inference over reference frames provides a unifying framework for both motion integration and segmentation (Bill et al., 2022, 2020; Gershman et al., 2016; Penaloza et al., 2024; Shivkumar et al., 2025). Causal inference is a Bayesian computational motif for inferring when to combine information across cues (Körding et al., 2007), and it has recently been proposed to be a universal computation across the cortex (Shams & Beierholm, 2022). This line of work suggested that the brain infers the causal structure underlying the elements moving in a visual scene, and uses that structure to determine the reference frames within which to represent the motion of each element. In this context, a reference frame is the coordinate system in which motion of objects is computed. For example, a passenger walking on a moving walkway moves slowly relative to the walkway but fast relative to the ground. In causal inference models of motion perception, each moving element is assigned to a reference frame, and its retinal velocity is the sum of the frame’s velocity and the element’s velocity relative to it. Reference frames can be nested, with the stationary world as the outermost frame. These models successfully explain behavior in psychophysical experiments using center-surround motion stimuli, effectively bridging the gap between normative computations and behavior (Bill et al., 2022, 2020; Gershman et al., 2016; Penaloza et al., 2024; Shivkumar et al., 2025).

In this study, we aim to bridge the remaining gap between normative computations in motion perception and neural activity. We built on a recently proposed normative Bayesian causal inference model of motion perception (Shivkumar et al., 2025), which accurately captures the perceptual effects of center-surround motion stimuli (Fig. 1a,d) using a biologically plausible architecture (Fig. 1b). In this model, participants’ motion-direction estimates were explained by inferring the reference frame in which center-surround motion stimuli were perceived. Here, we extended this model to the neural level: we derived its predictions for mean responses (tuning curves, Figs. 4 and 5, and Figs. S2 to S4) and neural variability (Fig. 5, and Figs. S2 and S3) across the full 4-dimensional stimulus space (direction and speed of center and surround motion). We compared these predictions with classic neural recording studies (Allman et al., 1985a; Born, 2000; Tanaka et al., 1986), which remain among the most detailed parametric characterizations of surround direction and speed effects in MT, yet still lack a unified account. Our model predictions captured a wide range of center-surround interaction effects (Figs. 5 to 7, and Figs. S2 to S5), and reflected both the representation of motion relative to a reference frame, inferred by causal inference, and the representation of motion in retinal coordinates. They are also compatible with a broad range of classic neurophysiological findings (Figs. 6 and 7, and Fig. S5), providing a normative explanation for why surround motion facilitates or suppresses neural responses, and when these effects are direction dependent. We focus on center-surround motion and area MT because the model is defined over velocities, and because MT is where center-surround motion interactions and basic tuning properties have been most extensively characterized in primates. However, the same logic, contextual modulation determined by an inferred grouping of scene elements, could be applied to other features and areas, as previous work has done for V1 (Coen-Cagli et al., 2012, 2015). Our results suggest future experiments – both to provide more comprehensive tests of the normative theory of Bayesian causal inference in motion perception and to link it to circuit-level mechanistic models.

**Figure 1:**
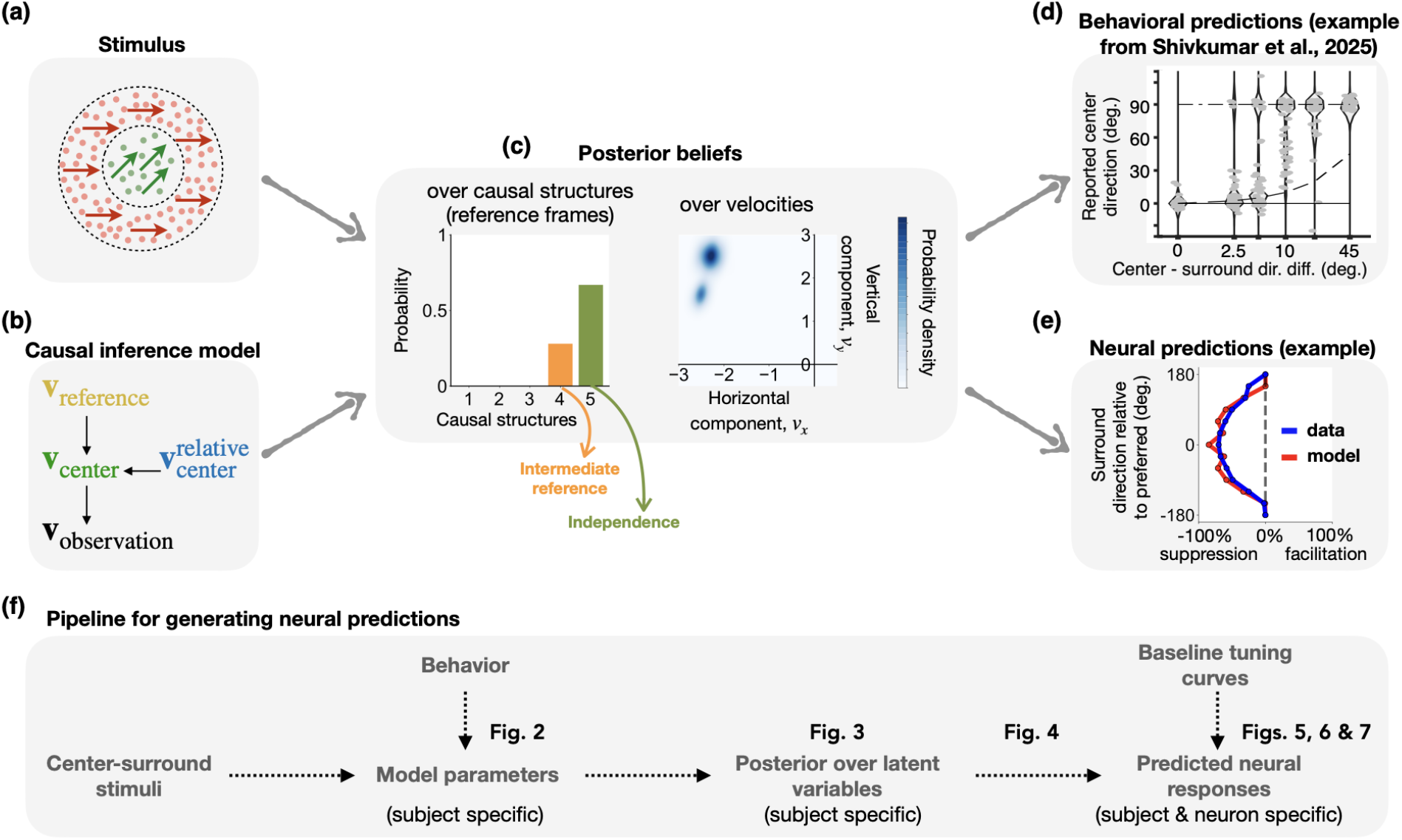
A normative, Bayesian framework for linking center-surround processing in perception and neural activity. This figure provides a conceptual overview of the study’s approach, from the experimental paradigm and Bayesian model to its connection with behavioral and neural data. **(a) Human experimental paradigm**. Observers view a central moving dot patch (green) within a moving surround (red) and report their perceived direction of the center. The retinal velocities of the center and surround are depicted by green and red arrows, respectively. **(b) Generative model for the observed stimulus velocity**. The model assumes the true retinal center velocity, **v**_center_, is equal to the vector sum of the reference frame velocity, **v**_reference_, and the center velocity relative to that reference frame, 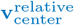. This true center velocity then produces a noisy sensory measurement, *v*_observation_. **(c) Posterior beliefs**. The brain infers posterior beliefs over latent variables, such as causal structure and velocities. The example shows posteriors where the observer infers two possible causal structures (on the left): Intermediate reference (perceiving motion relative to a reference frame intermediate between the center and surround motions) and Independence (perceiving the two patches as moving independently). The posterior over 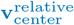 (on the right) reflects this, showing two modes that correspond to these two structures. **(d) Connecting the model to behavior**. The model is constrained by fitting it to human psychophysical data. The panel shows reports (gray dots) and the model’s posterior predictions (black violins) for an example observer from Shivkumar et al. (2025). The plot displays the observer’s perceived direction of the center stimulus (y-axis) as a function of the directional difference between the center and surround (x-axis). **(e) Connecting the model to neural activity**. The model’s posterior beliefs, constrained by behavior, are then used to predict the modulation (suppression/facilitation) of single MT neurons in response to complex surround motion. **(f) Overall research pipeline**. In summary, the framework uses a pipeline in which the model is first fitted to behavioral responses (Fig. 2). These subject-specific parameters are then used to compute posterior beliefs for any center-surround stimulus (Fig. 3). Finally, these posteriors are combined with baseline neural tuning curves (measured with a stationary surround) to generate testable predictions (Fig. 4) for how single neurons will respond to complex moving surrounds (Fig. 5).

**Figure 2:**
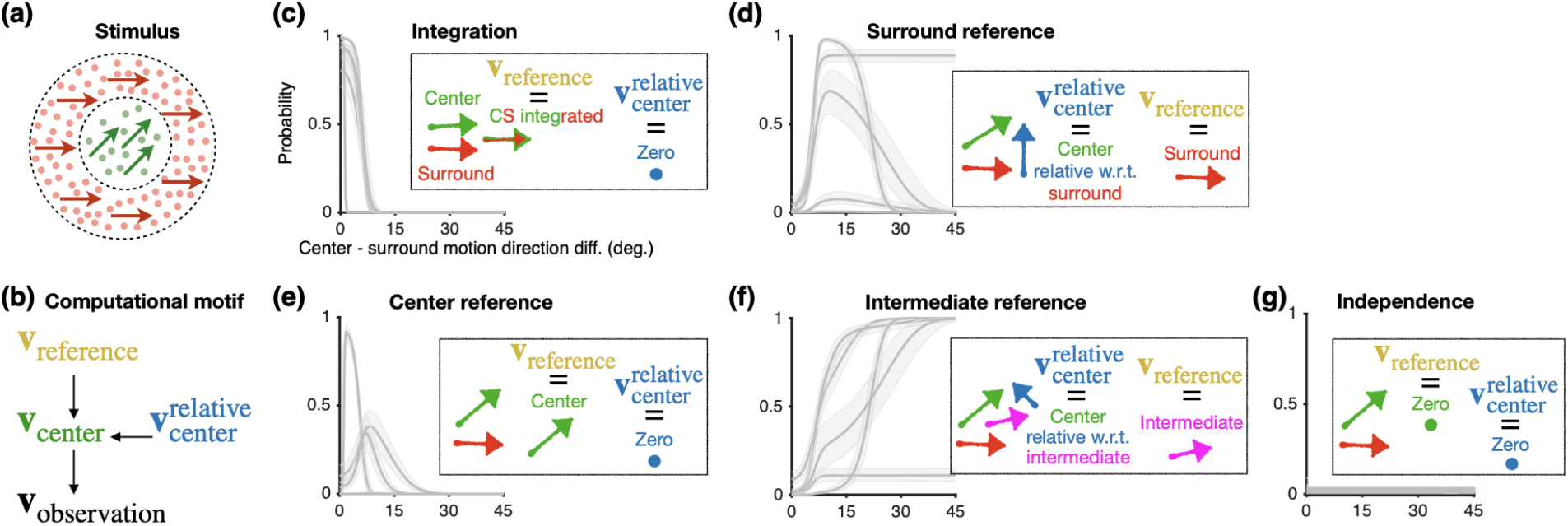
A hierarchical Bayesian model for inferring the causal structure of visual motion. **(a)** Same as Fig. 1a. **(b)** Same as Fig. 1b. **(c-g)** Posterior probability assigned to competing causal motion structures. The model considers 12 hypotheses for the relationship between center and surround motion. Here, we show 5 of the 12 structures depicted schematically (arrows: motion; dots: stationary) in the inset boxes. These include four structures assuming a common cause (c-f) and one assuming independent causes (g). The plots show the median posterior probability distributions for each of five different observers (solid grey lines), with 95% credible intervals (shading), as a function of the directional difference between the center and surround stimuli. Observers consistently assigned negligible probability to the independent-causes structure (g). Adapted from Shivkumar et al. (2025).

**Figure 3:**
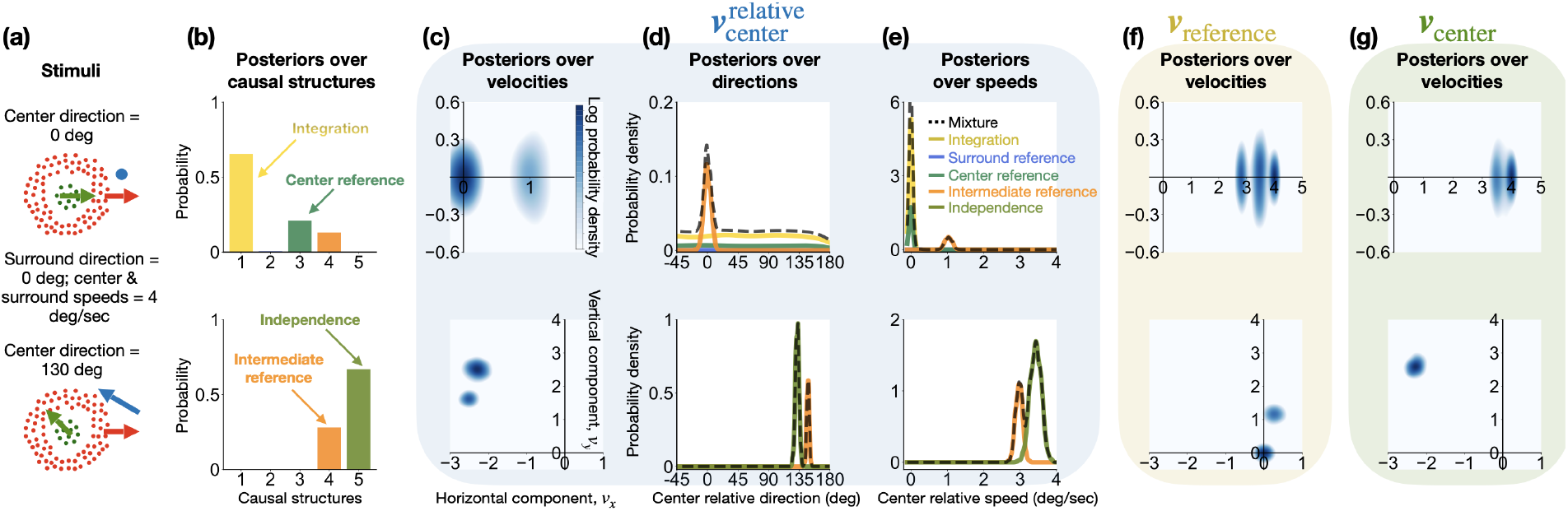
Causal inference generates complex, multi-modal posterior beliefs about latent motion variables. Posteriors were computed for observer #2 from Shivkumar et al. (2025) in response to two distinct center-surround stimuli (top vs. bottom row). **a:** The center-surround stimuli. The surround motion was fixed at 0° (red arrow), while the center motion differed between the two examples (green arrow). The relative velocity vector (center minus surround velocity) is shown in blue. The blue dot in the top row represents zero velocity. **b:** Posterior probabilities over the five (out of 12 total) causal motion structures (see Fig. 2c-g). Note how the most probable structure changes depending on the stimulus. **c-e:** Posterior beliefs about relative motion 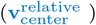. These panels show the full 2D posterior over velocity, and the marginal posteriors for direction and speed (dashed line: posterior beliefs; colored lines: individual mixture components corresponding to the motion structures). The multi-modal and complex shapes reflect the observer’s uncertainty over the causal structure. **f and g:** Posterior beliefs about the reference frame (**v**_reference_) and retinal motion (**v**_center_). Marginal posteriors for direction and speed are not shown for these variables. To aid visualization, all 2D velocity posteriors (c, f, g) are displayed using log probability density, which makes secondary modes visible. In contrast, the 1D marginal posteriors (d, e) are plotted using linear probability density.

**Figure 4:**
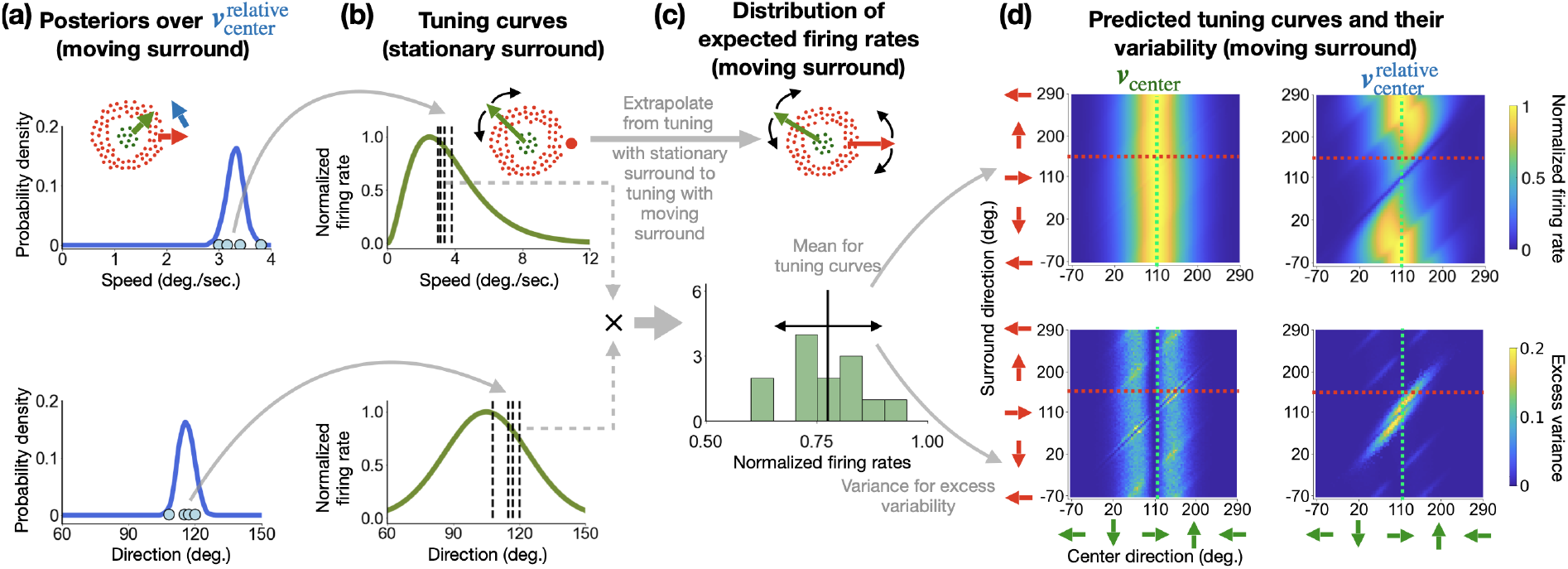
Linking posterior beliefs to neural activity via the neural sampling hypothesis. Schematic of the linking model used to generate neural predictions from the Bayesian observer model’s posterior distributions. All firing rates are normalized. **(a)** For a given center-surround stimulus (inset), the model’s posterior belief over a latent variable (here, 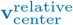) is approximated by a set of discrete samples (blue dots), in line with the neural sampling hypothesis (Fiser et al., 2010; Hoyer & Hyvärinen, 2002). **(b)** A baseline tuning curve is measured (here, in response to a stimulus with a stationary surround). This baseline serves as a look-up table to convert each sample from the test posterior (here, in response to a stimulus with a moving surround) into a predicted firing rate. **(c)** The mapped activities for speed and direction are multiplied (symbol ×), yielding a full distribution of predicted firing rates. The mean (solid black line) and variance (black arrow) of this distribution form our predictions for the neuron’s response to the specific stimulus in (a). **(d)** This entire process is repeated for a wide range of center-surround stimuli to generate complete 2D tuning curves. Shown are the predicted mean firing rate (top) and excess variance beyond the variability of a simple Poisson process (bottom) for a neuron assumed to represent either retinal motion (**v**_center_, left) or relative motion (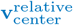, right). Model parameters are from observer #2 shown in Figs. 3 and 5.

**Figure 5:**
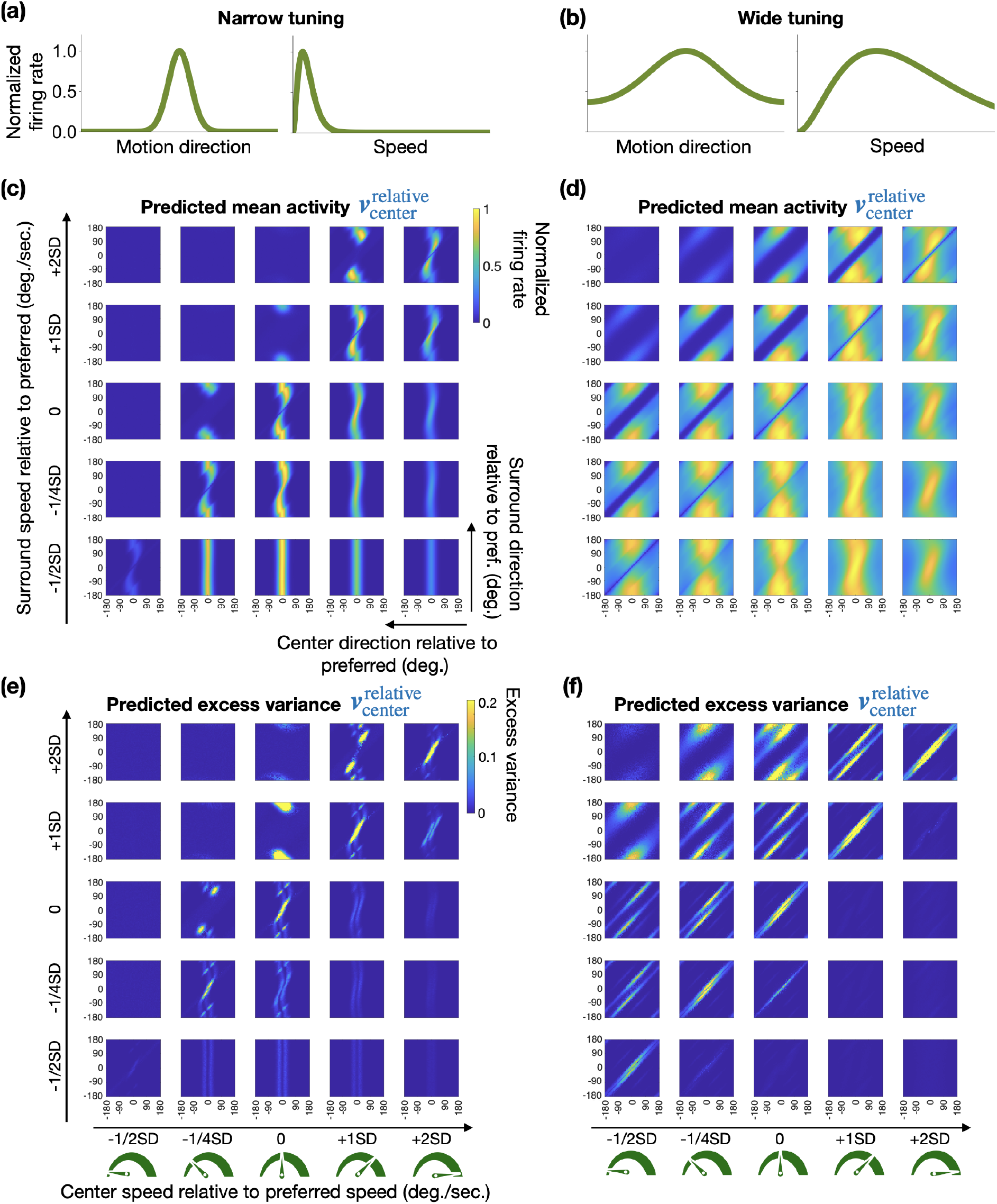
Predicted 4D tuning curves reveal complex signatures of causal inference. Predictions are based on parameters for observer #2 in Shivkumar et al. (2025) and two hypothetical neurons with different tuning curve shapes. **(a)-(b)** Schematics of the narrow and wide tuning curves used to generate predictions. **(c)-(d)** Predicted mean firing rates for model neurons that represent 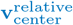, shown as a function of center direction (x-axis), surround direction (y-axis), center speed (columns), and surround speed (rows). Axes are relative to the neuron’s preferred direction and speed (where zero on both axes indicates the preferred direction). The key signature of causal inference is the complex pattern in the predicted tuning profile, which consists of a mixture of different diagonal interactions corresponding to the different reference frames under different motion structures. **(e)-(f)** Predicted excess variance beyond the variability of a simple Poisson process for the same conditions as (c-d). All firing rates are normalized.

**Figure 6:**
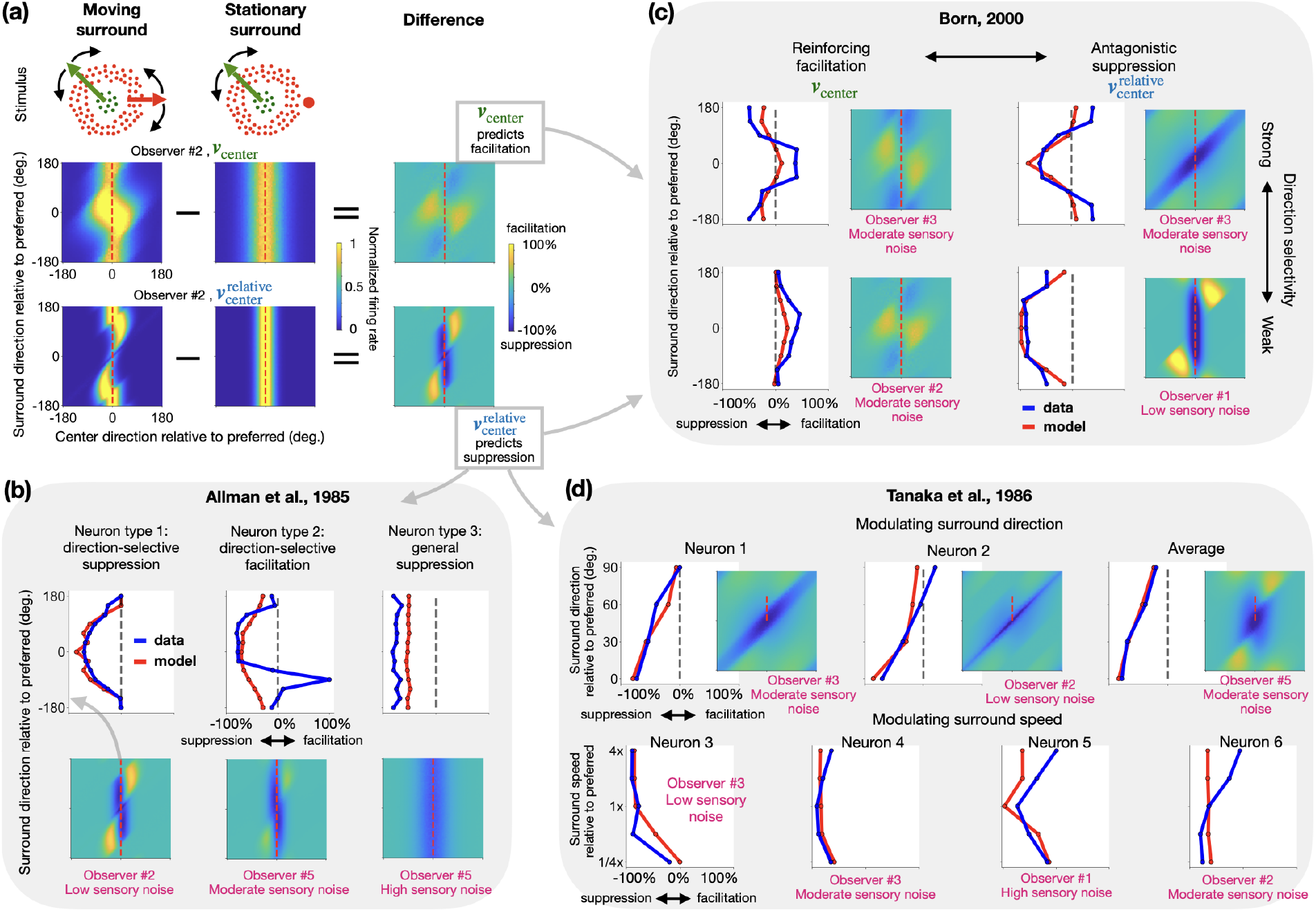
Model predictions qualitatively capture diverse surround modulation patterns in empirical MT data. The model’s predictions are compared to previously published single-neuron data from monkeys. In all heatmaps and plots, axes are expressed relative to the neuron’s preferred direction (labeled as zero). **(a)** Surround modulation is computed by subtracting the predicted response to a stationary surround (middle heatmap) from the response to a moving surround (left heatmap). The top inset illustrates the stationary- and moving-surround stimulus conditions. The model predicts strong suppression for neurons encoding 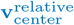 (bottom row, right heatmap, dark blue). This suppression is concentrated along the diagonal where center and surround move in similar directions, and along the vertical midline, where the center moves in the neuron’s preferred direction. The latter arises because with a moving surround, the center’s motion is represented relative to a reference frame between center and surround, which shifts the neuron’s peak response away from the preferred center direction (the bent band in the left heatmap). The response at the preferred direction therefore drops below its stationary-surround level. These neurons are also facilitated, but only when the center motion differs from the neuron’s preferred direction (bottom row, right heatmap, yellow regions). Neurons encoding **v**_center_ are only weakly modulated by surround stimuli (top row, right heatmap). Under moderate to high sensory uncertainty, however, they are weakly facilitated also when the center moves in the neuron’s preferred direction, i.e., along the experimentally tested cross-section (top row, right heatmap, at the horizontal midline; see also panel c, left column), consistent with facilitation observed empirically. The dashed red lines indicate the cross-section corresponding to the stimuli used in experimental studies. **(b)** Comparison to three modulation types from Allman et al. (1985a). We performed qualitative model fitting; see main text and Methods 4.7 for details. Model predictions assuming 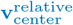 coding (red lines) are overlaid on average empirical data (blue lines). The plots (top row) show modulation (facilitation/suppression, x-axis) as a function of surround direction (y-axis), while the center patch moves in the neuron’s preferred direction. The full 2D predictions are shown as heatmaps (bottom row), with the dashed red line indicating the plotted cross-section, which corresponds to the experimental data we aim to match. Pink text indicates the best-matching observer model and sensory uncertainty level (“low,” “moderate,” and “high” correspond to the fitted noise value from Shivkumar et al. (2025), 10x, and 100x that value). **(c)** Comparison to four functional classes from Born (2000). The format is the same as in (b). The model accounts for the different neural classes by assuming different latent variable encodings. Predictions based on **v**_center_ under moderate uncertainty capture the observed facilitation effects (left column). In contrast, predictions based on 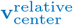 capture the observed suppression effects (right column). **(d)** Comparison to surround modulation from Tanaka et al. (1986). Following the same format, model predictions for 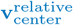 (red lines) are compared to data from example neurons and population averages for various surround conditions: in the top row, surround direction was varied while speed was fixed at the neuron’s preferred speed; in the bottom row, surround speed was varied while direction was fixed at the neuron’s preferred direction. The heatmaps and red dashed lines correspond to the direction manipulations in the top row. Corresponding heatmaps for the speed-varying data (bottom row) are not shown, as they would represent a different 3D parameter space that also contains surround speed.

**Figure 7:**
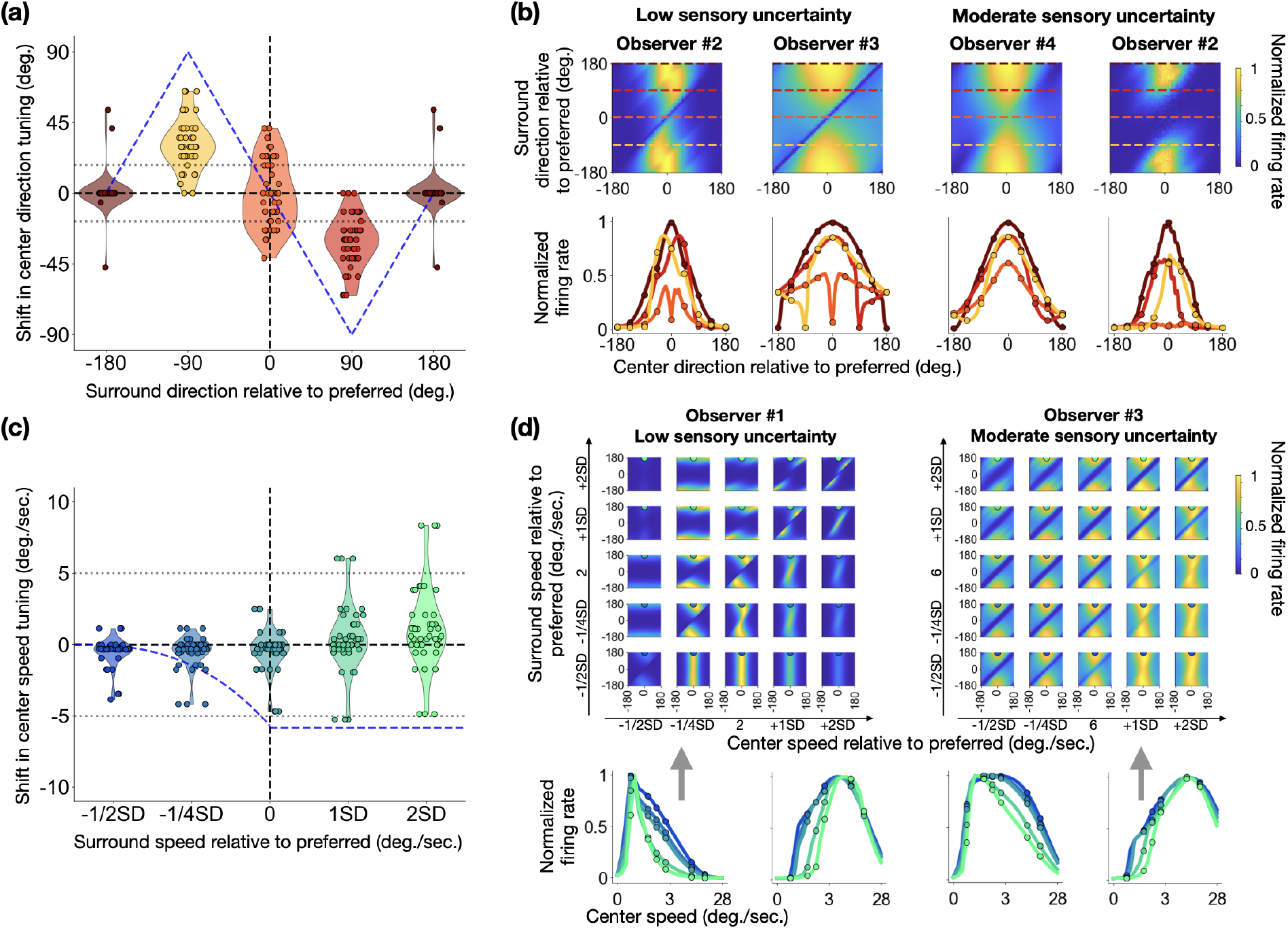
The model predicts when tuning curve shifts due to surround motion are large and when they are minimal. In all panels, axes are plotted relative to the neuron’s preferred direction and speed (labeled as zero). Specific points on the tuning curves correspond to the stimulus conditions tested in Born (2000). **(a)** Predicted shifts in center direction tuning (y-axis) as a function of four different surround directions (x-axis). Violin plots show the distribution of shifts across 45 combinations of observer parameters and neuron tuning profiles (5 observers × 9 tuning curves). The predicted shifts for some of the observers fall within the empirically observed range from Born (2000) (dotted horizontal lines), and most shifts are below those predicted by a pure surround-relative model under the surround reference structure (blue dashed line). **(b)** Example 2D tuning curves (top) and their 1D cross-sections (bottom) for direction. The colored dashed lines in the heatmaps indicate the surround directions for which the center tuning curves below are plotted. Predictions are shown for low (fitted values) and moderate (10x fitted values) levels of sensory uncertainty. The shift of the peaks is smaller for moderate sensory noise, consistent with the points around y=0 in (a). **(c)** Predicted shifts in center speed tuning, analogous to (a). The x-axis represents surround speed scaled by the tuning width (SD). The predicted shifts are smaller than what is predicted by a pure relative-motion model (blue dashed line) and mostly fall within the empirically measured range (dotted horizontal lines). **(d)** Example 4D tuning curves and 1D cross-sections for speed. The top row displays two examples of the predicted 4D tuning curves, showing firing rate as a function of center speed (columns) and surround speed (rows). The bottom row shows four example 1D center speed tuning curves. The first and last of these are cross-sections corresponding to the 4D predictions shown directly above them, while the two middle curves are additional examples.

## 2 Results

Our paper is structured as follows. First, we introduce the Bayesian causal inference model that we adapt, along with the center-surround motion experiment and the behavioral results used to fit the model parameters. Next, we examine the variety and complexity of posterior beliefs inferred by the model in center-surround motion tasks. We then describe how neural predictions – including tuning curves and response variability – are derived from the model. Following this, we characterize the center-surround interactions present in these predictions. Finally, we demonstrate that Bayesian causal inference qualitatively accounts for a wide range of neural center-surround interactions observed in MT.

### 2.1 Task, model, and prior empirical results

We used the model of Shivkumar et al. (2025) with parameters inferred from behavioral data of five human observers tested in their experiment (Fig. 2c-g). Participants reported their perceived motion direction of a central target patch composed of multiple moving dots (Fig. 2a, green dots), while the target was surrounded by a ring of dots that were either stationary (control condition) or moving (Fig. 2a, red dots; see Methods 4.1 for details on the stimulus). The experiment was designed to measure how participants’ motion direction estimates are influenced by the directions and speeds of both the center and the surround. This type of stimulus is commonly used to investigate center-surround processing in motion perception and in neurophysiology (e.g., Adelson and Bergen, 1985; Born, 2000; Tadin et al., 2008, 2019; Tanaka et al., 1986).

The hierarchical Bayesian causal inference model from Shivkumar et al. (2025) proposes that the brain jointly infers the appropriate reference frames (**v**_reference_ in Fig. 2) for each stimulus and its motion in that reference frame (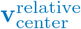 in Fig. 2). Formally, the retinal velocity of the center patch is the noisy sum of these two components, and the observer only has access to a noisy observation of it:

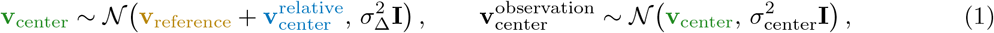

where *N* denotes a Gaussian distribution, 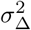 represents computational noise in the velocity computation, 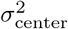 represents sensory noise, and **I** is the 2 × 2 identity matrix. Velocity variables, **v**, are two-dimensional vectors in Cartesian coordinates. The surround patch is described in the same way. Relative velocities have a mixture prior, 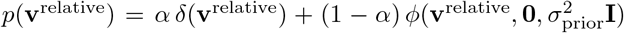: the delta component expresses the expectation that elements are often stationary relative to their reference frame, and *ϕ* a Gaussian probability density, with zero mean and variance 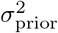, which embodies the assumption that velocities are generally small. The mixture weight *ε* is the prior probability that an element is stationary relative to its reference frame (see Methods 4.2).

All prior and noise parameters were fitted jointly to each observer’s direction reports (Shivkumar et al., 2025). The experimental design makes them identifiable, and the joint posterior over parameters shows how well they are constrained (see Methods 4.2).

Applied to this center-surround stimulus (Fig. 2a), the model assumes that observers infer the probability of 12 possible causal structures, *s*, each specifying whether the center and surround share a reference frame and which velocities are stationary within their reference frame. Because the observer does not know which structure generated the stimulus, the posterior over each latent velocity is a mixture of the posteriors under each structure, weighted by the posterior probability of that structure:

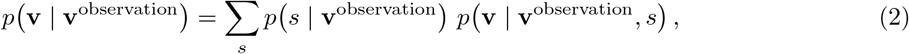

where **v** stands for any of the latent velocities (**v**_center_, **v**_reference_, or 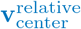), and 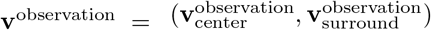 denotes the noisy observations of the center and surround velocities. Multi-modal posteriors can therefore arise whenever more than one causal structure is plausible.

Notably, in Shivkumar et al. (2025), four of these structures had significant probability mass. While we use these four for illustration, our model’s predictions incorporate all 12 structures (i.e., the model always marginalizes over all 12 structures, both when fitted to behavior and when generating neural predictions; no structure is excluded from the computation). In addition to these four, we also highlight a fifth causal structure (also one of the 12) in which the center and surround move independently. In this “independence” structure, motion is perceived relative to a world reference frame (Fig. 2g and Fig. S1k). This can be considered as motion segmentation with respect to a world reference^1^. This structure had negligible mass in fits to data performed by Shivkumar et al. (2025), presumably due to the patches’ proximity, shared temporal dynamics, and small motion differences. However, the wider stimulus range (relative to those used in the original experiment) that we explore here makes the independence structure a more plausible alternative.

The first of the four main causal structures is classical motion “integration” (Braddick, 1993; Tadin & Lappin, 2005; Tadin et al., 2003) (see Fig. 2c; generative model in Fig. S1c). Observers infer this structure as the most probable when the difference in motion between center and surround patches is small, leading to a “cue-combined” perception in which both motions are integrated (Fig. 2c, curves). In contrast, motion segmentation (Braddick, 1993; Tadin et al., 2019), the second causal structure, arises when the motion differences between center and surround are substantial. We term this second structure “surround reference” (Fig. 2d and Fig. S1e) because observers segment the center’s motion from the surround, perceiving it relative to the surround’s motion. Interestingly, this structure was dominant for only one of five observers, playing a less significant role for the other participants (Fig. 2d).

The third and fourth causal structures account for behavioral variability beyond canonical integration and segmentation. In the third structure, observers perceive the surround motion relative to the center group, using the center as a reference frame (Fig. 2e and Fig. S1g). Since observers in this task report the motion direction of the center patch, this “center reference” structure generates responses that are unaffected by the surround patch, much like the independence structure (Fig. 2g, Fig. S1k). The fourth structure captures when observers adopt an “intermediate reference frame” between the center and surround patches’ motions, perceiving motions of both patches relative to this intermediate frame. Most observers put significant probability mass on this structure across many different center-surround stimulus configurations (Fig. 2f and Fig. S1i).

These latter two structures can be easily miscategorized as classical integration or segmentation. For instance, the center reference structure can be confused with motion integration when center and surround motions are very similar, because the theoretically integrated percept would lie very close to the center’s actual motion. Similarly, the intermediate reference structure can be mistaken for motion segmentation if one assumes that the subtraction of the surround’s velocity from the center’s is incomplete (Shivkumar et al., 2025). The Bayesian causal inference model resolves these ambiguities, differentiating between all of these structures and explaining diverse behavioral phenomena through a single computational principle.

### 2.2 Causal inference produces complex posterior beliefs

In order to generate neural predictions for arbitrary center-surround stimuli, we independently varied center-surround velocities (each consisting of direction and speed) and computed the posteriors over all latent variables in the Bayesian causal inference model using parameters fitted to the five observers in Shivkumar et al. (2025). The posteriors can be computed using any parameter values hypothesized by the researcher. However, the parameters fitted to real behavioral responses will reveal posterior beliefs that are backed up by empirical data. The model (Fig. 2b) contains three latent variables that are involved in representing the center patch’s motion: **v**_reference_, **v**_center_, and 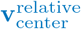. **v**_reference_ denotes the motion of the reference frame in retinal coordinates. **v**_center_ represents the motion in retinal coordinates while 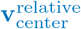 represents the motion relative to the reference frame. Note that these latent variables are distinct from the model’s perceptual read-out. The reported percept is a separate, structure-dependent variable: the non-zero relative velocity that is lowest in the hierarchy (Methods 4.2, Eq. 11, Shivkumar et al., 2025). Our neural predictions concern the posteriors over the latent velocity variables themselves, not over the percept.

As a general principle, the posterior over **v**_center_ is only minimally affected by surround motion, typically forming a simple, Gaussian-like distribution centered on the physical velocity of the center patch, reflecting the uncertainty due to observation noise (Fig. 3g, bottom). In contrast, the posteriors for **v**_reference_ and 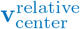 are strongly modulated by the surround, often displaying complex, multi-modal forms characteristic of causal inference (Fig. 3c-f). In the main text, we illustrate our findings using a representative observer (observer #2 in Exp. 1 of Shivkumar et al. (2025); Figs. 3 to 5), with the results for all participants available in Figs. S1 to S3 and S5.

To illustrate the complexity arising from the causal inference process, we first examine the model’s posterior beliefs for a nominally simple stimulus where the center and surround move identically (Fig. 3, top row). Contrary to expectation for such a simple stimulus, most observers did not infer a single causal structure. Instead, they assigned significant probability mass to a mixture of competing structures. For instance, observer #2 in Shivkumar et al. (2025) assigned high probability to three structures: motion integration, the center patch acting as a reference frame, and an intermediate reference frame. This mixture of inferred structures directly shapes the posterior distributions. The posterior over the reference frame (**v**_reference_ ; Fig. 3f) consequently shows three distinct peaks corresponding to these probable structures. The dominant peak aligns with motion integration (around 0° direction and 4 deg/s speed), while secondary modes correspond to the center-reference (middle peak at 3.7 deg/s) and intermediate-reference (left peak at 2.8 deg/s) interpretations. Notably, the peak for the center-reference mode reflects a slower motion than the true stimulus speed; this discrepancy is a direct consequence of the slow-speed prior inherent in the model (Shivkumar et al., 2025).

Correspondingly, the posterior over relative velocity (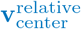 ; Fig. 3c-e) is not a simple peak at zero. While its main component is concentrated at zero velocity (corresponding to both integration and center-reference structures), it exhibits one additional mode that reflects the observer’s belief in the intermediate-reference structure. This additional mode is visible as a smaller peak in the full posterior view (top, blue contour around *v*_*x*_ = 1 deg/s in Fig. 3c). This single mode manifests in the marginal distributions as both a peak at 0° for direction (in addition to the uniform distribution representing the stationary component; Fig. 3d) and a secondary peak near 1 deg/s for speed (Fig. 3e).

The posterior over retinal motion (**v**_center_ ; Fig. 3g), while typically unimodal, also displays a subtle bimodality in this case. This corresponds to the center-reference structure, which, as noted, was inferred to be slightly slower than the true stimulus speed. This could indicate a small perceptual bias for this observer even under this simple center-surround stimulus. Together, this example demonstrates that due to uncertainty over the underlying causal structure, the model’s posterior beliefs are highly complex even when the stimulus itself seems unambiguous to the experimenter.

A second example highlights how the model arbitrates between competing causal structures when the directional difference between the center and surround motion is large (Fig. 3, bottom row). The same example observer infers two probable structures for this center-surround stimulus (Fig. 3b): (1) that the motions are independent, or (2) that they share a common intermediate reference frame. Each of these interpretations corresponds to a distinct mode in the bimodal posteriors for **v**_reference_ and 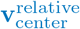. The independent-motion structure generates a mode for **v**_reference_ at zero velocity (Fig. 3f), with a corresponding mode for 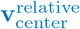 at the retinal velocity of the center patch (top blue contour in Fig. 3c). Conversely, the intermediate-reference structure produces a second mode for **v**_reference_ at the inferred reference velocity (bottom blue contour around *v*_*y*_ = 1 deg/s in Fig. 3f), with 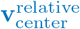 peaking at the center’s velocity relative to this moving frame (bottom blue contour in Fig. 3c). Note that this stimulus, with a large direction difference between center and surround, lies outside the range tested in the experiments of Shivkumar et al. (2025), where the independence structure received negligible probability. Its high probability here is therefore an out-of-distribution prediction of the model.

### 2.3 A novel sampling-based method for predicting single neural activity from posterior beliefs

To bridge the gap between our normative, Bayesian causal inference model and neural data, we introduce a novel linking method which posits that single-neuron activity reflects samples from the model’s posterior beliefs (i.e., the neural sampling hypothesis; Fiser et al., 2010; Hoyer and Hyvärinen, 2002). Neural sampling enables us to generate neural predictions that reflect the complex, multi-modal nature of the model posteriors. Our method provides a general framework for predicting single neural responses to any “test” stimulus from any Bayesian model (Fig. 4). Specifically, it posits that the response to the test stimulus can be derived from the same neuron’s tuning curve measured for a simpler “baseline” stimulus set. This baseline tuning curve acts as a “look-up table”: each sample from the test posterior is converted into an individual firing rate (using the look-up table), and aggregating all these rates forms a full distribution of predicted responses (more details are provided in the Methods 4.4). This method can be applied in any domain, as long as its two key components are well-defined: (1) the Bayesian model parameters to compute posterior beliefs and (2) the tuning curve parameters to link posterior samples to firing rates.

To apply this method to our center-surround motion stimuli, we define these two components as follows. For the Bayesian model parameters, we used the unique parameters fitted to each of the five human observers in Shivkumar et al. (2025). For the tuning curve parameters, we constructed tuning profiles approximating the 25th, 50th, and 75th percentiles of empirically measured MT tuning curves from monkeys (DeAngelis & Uka, 2003). While this combines human behavioral models with monkey neurophysiology, it provides a strong proof of principle. Ideally, both behavioral and neural data would be collected from the same subjects.

We then implemented the linking procedure (Fig. 4). We assume that each neuron represents, and hence only depends on, the posterior over a single latent variable, either the retinal velocity of the center (**v**_center_) or its velocity relative to the inferred reference frame 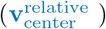. First, for a given test stimulus (moving surround), we generated samples from the model’s posterior distribution over this latent variable (Fig. 4a). Second, we used the baseline tuning curve (derived from the parameters of MT neurons’ tuning curves measured with a stationary surround in DeAngelis and Uka, 2003) as a look-up table (Fig. 4b) to map each posterior sample to a predicted firing rate yielding a full distribution of predicted firing rates (Fig. 4c). Finally, we calculated the mean and variance of this distribution, which serve as our predictions for the neuron’s mean firing rate and response variability (Fig. 4d). Formally, for a neuron with center-only (i.e., with stationary surround) tuning curve *f* (**v**) = *f*_speed_(*u*) *f*_direction_(*θ*) (where *u* and *θ* are the speed and direction), the predicted mean response and variability are

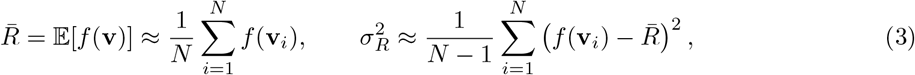

where **v**_*i*_ are samples from the posterior over the latent variable encoded by the neuron 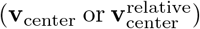.

In essence, this method leverages a neuron’s simple motion tuning curve to predict its complex response to center motion with moving surround. It achieves this by assuming the neuron’s firing is determined not by the raw stimulus, but by the brain’s internal belief about the probable motion components and their reference frames, as represented by the samples from the model’s posteriors (full details of the method are provided in Methods 4.4).

### 2.4 Predicted 4D tuning curves reveal complex signatures of causal inference

Applying our linking method, we generated predictions for neurons representing posterior beliefs over either **v**_center_ or 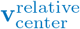 within their classical receptive field, matching the empirical studies we aim to explain (Allman et al., 1985a; Born, 2000; Born & Bradley, 2005; Born & Tootell, 1992; Tanaka et al., 1986). In contrast, the spatial extent of a receptive field representing **v**_reference_ is more ambiguous, and we therefore do not analyze predictions for **v**_reference_ here.

First, we tested the hypothesis that two classes of MT neurons (Born & Bradley, 2005) – those insensitive to the surround and those with strong surround modulation – correspond to neurons encoding **v**_center_ and 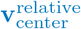, respectively. We generated four-dimensional (4D) tuning curves (Fig. 5, and Figs. S2 and S3) as a function of center direction (x-axis in each panel), surround direction (y-axis in each panel), center speed (increasing across columns), and surround speed (increasing across rows). In each panel, a neuron whose responses are only determined by the center stimulus would produce vertical bands of activity. Consistent with our hypothesis, the predictions for a neuron encoding retinocentric motion (**v**_center_) show such uniform vertical bands across all conditions (Fig. 4d, top-left panel, and Fig. S3), and thus align with the surround-insensitive neuron class. In stark contrast, predictions for a neuron encoding relative motion 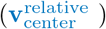 are substantially modulated by the surround, revealing complex center-surround interactions that match the surround-dependent neuron class (Fig. 4d, top-right panel and Fig. S2). We will now use the neural predictions of 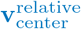for the representative observer #2 in Shivkumar et al. (2025) to explain the key features of the causal inference model’s predictions below (predictions for **v**_center_ and for the remaining four observers shown in Figs. S2 and S3).

To understand the pattern in Fig. 4c, d, note that a neuron representing relative motion, 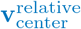, responds to the velocity difference between the center and the inferred reference frame. When the surround serves as this reference frame, the velocity difference is smallest where center and surround move in the same direction. The response is therefore organized along the diagonal of each panel, not vertical or horizontal bands. We call this a diagonal interaction and quantify it by the loss of symmetry (Methods 4.6). Fig. S4 illustrates the range of diagonal interactions predicted by the model across a broad sample of parameters around the fitted values, ordered from the strongest to the weakest diagonal interaction according to this measure. Importantly, such a diagonal interaction is not separable: it cannot be written as a product or sum of a center-direction tuning and a surround-direction tuning, as would result from a fixed surround gain. It therefore cannot be explained by surround modulation that depends on the surround direction alone.

The mechanism for this diagonal structure is twofold. First, the observer’s posterior over 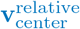 peaks at the velocity difference between the center and the inferred reference frame. Second, the predicted firing rate is highest when these posterior peaks align with the neuron’s preferred direction and speed. Consequently, the more an observer interprets the surround as a reference frame, the stronger this diagonal interaction becomes (see the strong diagonal structure of a pure surround-relative representation for observer #1 in Fig. S2), while a motion representation relative to an intermediate reference frame results in a weaker diagonal interaction (e.g., observer #2 in Fig. 5). The strongest diagonal interaction is therefore predicted under classical motion segmentation, a perceptual outcome in which the surround is treated as the sole reference frame for the center’s motion (Braddick, 1993; Tadin & Lappin, 2005).

It is important to note that all predictions in Fig. 5 (and Figs. S2 and S3) are displayed relative to the neuron’s preferred speed and direction, making these 4D tuning curves generalizable to any neuron with a similar tuning profile. The specific patterns of center-surround interaction, however, depend on two key factors. First is the neuron’s intrinsic tuning shape, with narrower and broader tuning curves yielding qualitatively different interactions (Fig. 5, left vs. right columns). Second is the observer’s perception, as variations in the fitted model parameters lead to substantial differences in the predicted responses across individuals (compare panels in Figs. S2 to S4). Thus, for a given observer and specific tuning curves in response to center motion alone (i.e., with stationary surround), each latent variable predicts exactly one 4D tuning curve, which makes these predictions directly testable.

Importantly, causal inference across potential reference frames shapes the diagonal interaction. When an observer infers a mixture of probable motion structures, the model’s posterior becomes multimodal (Fig. 3). This mixing of beliefs translates into complex response patterns. For instance, in Fig. 4d (top-right panel), the neural prediction for observer #2 is shaped by a mixture of three stimulus-dependent interpretations: (1) for large center-surround direction differences (beyond approximately 100°), the patches are inferred as independent, so that over this range of surround directions the response depends only on the center direction, producing vertical band segments confined to the top and bottom of the panel rather than bands spanning the whole panel; (2) for moderate differences, an intermediate reference frame is used, creating a curved peak near the diagonal; and (3) for similar directions, integration occurs (i.e., relative motion, 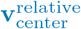, is zero), suppressing activity along the diagonal.

A second key characteristic of our Bayesian models is the assumption that observers not only form beliefs about motion but also represent the uncertainty in those beliefs. Under the neural sampling hypothesis (Fiser et al., 2010; Hoyer & Hyvärinen, 2002), this uncertainty is reflected in trial-to-trial variability: on each trial, a neuron’s firing rate corresponds to a sample from the posterior. We assume that spikes are then generated by a Poisson process with this fluctuating rate. The variance of the predicted rates (Eq. 14) therefore corresponds to variability in excess of Poisson noise (cf. Goris et al., 2014): the total spike-count variance is the mean rate (Poisson component) plus this excess term (the variance of the mean). Unlike in gain-modulation models, where the excess variance scales with the squared mean (Goris et al., 2014), here it reflects the uncertainty of the posterior and therefore has its own stimulus dependence, not tied to the mean rate.

These variance predictions are important because, unlike the mean-rate predictions, which hold for any linear distributional code (Methods 4.5), they are specific to sampling-based codes, and can therefore discriminate between encoding hypotheses that mean responses cannot distinguish.

We computed this predicted excess variance (i.e., the variance of the firing rates across posterior samples, Eq. 14) over the same four dimensions (center-surround directions and speeds) and found it also exhibits diagonal structures (Fig. 4d, bottom panels, Fig. 5e & f, and Figs. S2 and S3, bottom rows). This diagonal structure appears because the highest uncertainty in posterior beliefs – and thus the highest predicted firing rate variance – occurs when posteriors are multimodal. This happens when an observer considers multiple causal structures to be simultaneously plausible. For example, when center-surround directions are similar, both integration and segmentation with respect to an intermediate reference are plausible (see the high variability regions along the diagonal in Fig. 5e & f). Or when center and surround directions are far apart, making both independence and segmentation plausible (see the regions further away from the diagonal in Fig. 5e & f).

Crucially, high uncertainty in the latent variables at the neural level does not necessarily imply high behavioral variability as the final behavioral percept in the model is derived after integrating over all latent uncertainty, often resulting in a unimodal predictive posterior for decision (Shivkumar et al., 2025).

### 2.5 Comparing causal inference predictions to the empirical literature

The role of area MT in encoding motion remains a topic of debate. While some neurons were found to represent retinal motion and are unaffected by surround motion (Inaba et al., 2011, 2007; Newsome et al., 1988), others appear to encode motion relative to the surround (Allman et al., 1985a; Born & Tootell, 1992; Huang et al., 2007, 2008; Tanaka et al., 1986; Tzvetanov & Womelsdorf, 2008). Complicating this picture, studies have found a wide variety of additional center-surround interactions, such as direction-selective suppression, non-selective suppression, and even direction-selective and non-selective facilitation (Allman et al., 1985a; Born, 2000; Born & Bradley, 2005; Tanaka et al., 1986). No single theory yet captures all of these diverse center-surround effects (Born & Bradley, 2005). In this section, we show that our causal inference model predicts this observed diversity reasonably well. These varied response patterns emerge naturally from the model, depending on whether a given neuron encodes the retinocentric (**v**_center_) or relative-motion 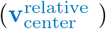 variable, and on the inferred motion causal structure determining the reference frame in which motion is perceived.

We qualitatively compared our model’s predictions against published data from three key neurophysiology studies: Allman et al. (1985a) (61 single units, 3 owl monkeys), Tanaka et al. (1986) (105 single units, 4 Japanese monkeys), and Born (2000) (169 single units, 18 owl monkeys). These studies used the same basic center-surround design: a center stimulus within the classical receptive field, moving in the neuron’s preferred direction and speed, and an independently moving surround, with a stationary or absent surround as a baseline condition. The studies differed in stimulus details, such as the sampled surround directions and speeds and the type of center stimulus (Methods 4.7).

A key challenge in this comparison is that the model’s predictions depend on both observer-specific model parameters and neuron-specific tuning curve parameters. Ideally, one would fit the model to an animal’s behavior and use tuning curves recorded from the same animal. Since this was not possible, we generated a diverse set of predictions using the five sets of model parameters from the human observers in Shivkumar et al. (2025) and a library of nine hypothetical but plausible tuning curves (Fig. S5a). This library was constructed by combining three distinct tuning widths for both speed and direction (3 speed widths × 3 direction widths), approximating the 25th, 50th, and 75th percentiles of MT tuning curve parameters measured by DeAngelis and Uka (2003). To evaluate whether these observed MT responses support our model, we compared this large set of predictions against the empirically measured responses. We first categorized the recorded neurons by their functional type of center-surround interaction. This grouping allowed us to assess whether each distinct neural response pattern could be explained by the causal inference framework. For each empirical center-surround interaction type, we then identified the best-matching prediction from our library (i.e., the one with the lowest root mean squared error; more details are provided in Methods 4.7).

We restrict these comparisons to mean responses because we are not aware of any study that reports trial-to-trial variability in MT as a function of surround motion. Existing analyses of MT variability vary other stimulus dimensions, such as the motion direction or texture of a stimulus confined to the receptive field (Gharaei et al., 2013; Ponce-Alvarez et al., 2013). The variability predictions therefore remain to be tested in future experiments.

#### 2.5.1 Neurons unaffected by surround motion

A subset of MT neurons exhibits no measurable modulation by surround motion. This response profile (which 8% of MT neurons show in Allman et al., 1985a) indicates that these cells respond selectively to center motion while remaining unaffected by motion in the surround. This response profile aligns with classical accounts of MT as representing motion in retinal coordinates (Inaba et al., 2011, 2007; Newsome et al., 1988). This pattern is predicted by the causal inference model for neurons representing the retinocentric latent motion variable (**v**_center_) as the posterior over this variable is largely unaffected by surround motion (Fig. 4d, left column, and Fig. S3).

#### 2.5.2 Neurons exhibiting direction-selective antagonistic suppression

The most common class of MT neurons exhibits strong suppression when the center and surround move in the same direction, with suppression decreasing as their directions diverge (Allman et al., 1985a; Born, 2000; Tanaka et al., 1986). The causal inference model predicts this suppression for neurons representing the relative velocity latent variable 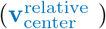. Mechanistically, this suppression arises because the model for relative motion 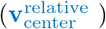 predicts a diagonal center-surround interaction for the moving surround condition, where activity is lowest along the diagonal (where the relative velocity is zero). When the response to a stationary surround is subtracted, this results in strong, direction-selective suppression (Fig. 6a, bottom row). In our model, this pattern emerges across all observer parameters and tuning curve shapes (Fig. S4, Fig. S5b), and the model’s predictions closely align with the measured neural data from multiple studies (Fig. 6b, left; Fig. 6c, right column; Fig. 6d, top). The strength of this center-surround effect depends on how closely the inferred reference frame aligns with the surround motion. For example, a surround reference causal motion structure (see Fig. 2d) produces the strongest diagonal center-surround interaction in the neural predictions.

However, previous studies could not measure the reference frame an observer inferred during viewing of a center-surround stimulus. This unmeasured variable is likely a substantial source of the variability observed across prior studies. Consequently, although a diagonal interaction is a key signature of relative motion, the actual neural data will deviate from the naive expectation of a pure diagonal center-surround interaction due to the complex relationship between the posterior and the stimulus via Bayesian causal inference over reference frames.

Moreover, the single cross-section of the 2D tuning space measured in most studies (e.g., Fig. 6, red cross-sections) is insuffcient to distinguish a true diagonal interaction – the key signature of relative motion – from other forms of surround suppression. Verifying such a representation would require testing multiple center and surround motion directions in combination.

Note that this suppression does not imply a segmentation percept. When the center and surround move together in the same direction, the observer most likely integrates them, so the inferred relative velocity of the center is close to zero, and a neuron representing the relative velocity, 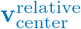 is suppressed. Thus, our model does not predict a one-to-one correspondence between perceptual states (integration or segmentation) and surround modulation.

#### 2.5.3 Neurons showing non-directional surround suppression

Approximately 30% of MT neurons in Allman et al. (1985a) (and a smaller fraction in other studies) exhibited suppression from the surround regardless of its direction (Born, 2000; Tanaka et al., 1986). In our model, such tuning arises for a neuron encoding 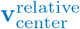 when the observer’s sensory uncertainty is moderate or high (Fig. 6b, right). High uncertainty broadens the posterior distributions over 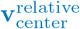, which averages out directional effects and leads to a more uniform, direction-independent suppression, consistent with these neurons’ responses. In this context, “low”, “moderate”, and “high” uncertainty correspond to the sensory uncertainty fitted to behavioral data in Shivkumar et al. (2025), and to levels 10x and 100x greater than that fitted value, respectively. These elevated values are justifiable because motion perception thresholds are typically lower in humans than in monkeys (Cherian et al., 2025; Law & Gold, 2008; Rangotis et al., 2024), and single-neuron responses presumably reflect greater uncertainty than behavioral output that integrates information from many neurons.

#### 2.5.4 Neurons with reinforcing surround modulation

Our model can also explain the “reinforcing modulation” observed in a subset of MT neurons, which constitutes a particular challenge for normalization-based models. For those neurons, responses increase when the surround moves in the same direction as the center (Born, 2000). This facilitation is predicted for two of our five observer models under specific conditions: that the neuron encodes the retinocentric variable (**v**_center_) and that sensory uncertainty is moderate (10x the value inferred from human behavior) (Fig. 6c, left column). When center and surround move with similar velocities, integration is the most probable causal structure. Integrating the two observations reduces uncertainty, thus the posterior over **v**_center_ becomes narrower than with a stationary surround, where the two patches are not fused. Because the predicted firing rate averages the neuron’s tuning curve over the posterior, a narrower posterior centered on the preferred velocity yields a higher mean rate, i.e., facilitation. This also explains why facilitation requires moderate sensory uncertainty: when the posterior is already much narrower than the tuning curve (i.e., low uncertainty), further narrowing has little effect.

Hence, the same stimulus that suppresses neurons representing 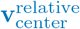 (Section 2.5.2) can facilitate neurons representing **v**_center_ . Moreover, a single neuron can switch between suppression and facilitation depending on the stimulus: neurons representing 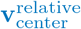 are facilitated for other combinations of center and surround directions (Fig. 6c, right column, yellow regions). With a stationary surround, the relative velocity of the center equals its retinal velocity. With a moving surround, the center’s velocity is instead represented relative to the surround or to an intermediate reference frame. When the center does not move in the neuron’s preferred direction, subtracting the reference velocity can bring the relative velocity closer to the preferred velocity, therefore the response increases relative to the stationary-surround condition (e.g., a cross-section 45° away from the preferred direction in Fig. 6a, bottom row). Previous studies, however, measured only one cross-section through this space, with the center moving in the neuron’s preferred direction (red dashed lines in Fig. 6). Along this cross-section, 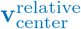 predicts suppression or no modulation, whereas facilitation is predicted only for **v**_center_ .

#### 2.5.5 Neurons showing surround facilitation orthogonal to their preferred direction

It might seem that our model is capable of explaining all observed phenomena, potentially rendering it unfalsifiable. We stress, however, that this is not the case. Our framework is constrained both by model parameter values fitted to behavioral data and by measured baseline tuning curves. Consequently, the model can only account for a small, specific subset of all possible four-dimensional center-surround tuning curves, which makes the agreement with the data described in previous sections a non-trivial success. This point is illustrated by the fact that a small fraction of neurons (8%) in Allman et al. (1985a) were facilitated by surround motion orthogonal to their preferred direction (Fig. 6b, middle column). Our model does not reproduce this effect under any parameter settings, suggesting these neurons may perform a different computation than the one described by our framework. It is noteworthy, however, that this specific finding involved a small proportion of neurons and lacked replication in later studies (Born, 2000; Born & Tootell, 1992; Tanaka et al., 1986).

#### 2.5.6 Neurons showing speed-dependent suppression

Tanaka et al. (1986) reported that surround suppression in MT is modulated by surround speed in addition to direction. Our model accounts for the main pattern they observed at the population level, stronger suppression at higher surround speeds, as illustrated by neuron 3 (Fig. 6d). This effect is robustly predicted for a neuron encoding 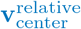, as its activity decreases monotonically with increasing surround speed across most parameter settings (Fig. S5c). Consequently, the model does not account for non-monotonic speed dependence, such as the C-shaped profile of neuron 4, nor for the opposite pattern – stronger suppression at lower surround speeds – which was observed in two other neurons from the same study (Fig. 6d, neurons 5 and 6). However, we are not aware of other studies that have reported this opposite effect.

#### 2.5.7 Neurons with minimal shifts in tuning under surround manipulation

To directly test whether neurons encode relative motion with respect to the surround (i.e., surround reference causal motion structure), Born (2000) recorded from neurons while jointly varying center and surround direction or speed. If neurons truly represent center motion relative to the surround, we would expect the center tuning to shift accordingly. However, the authors found little to no tuning curve shift, merely an overall response modulation. While this finding seems to challenge pure relative-motion models, the small number of neurons in these experiments (3 for direction, 9 for speed) and sparse stimulus sampling limit strong conclusions.

Our causal inference model demonstrates that a neuron representing relative motion (i.e., 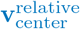) can indeed produce a response pattern similar to those observed by Born (2000). Moreover, our model provides specific predictions for when a neuron’s tuning shifts should be large and when they should be minimal or even zero, thereby accounting for the seemingly contradictory findings in the literature. In the model, the magnitude of the shift depends on the inferred reference frame. When the model infers an intermediate reference frame lying between the center and surround, it predicts minimal shifts with direction-independent response modulation observed by Born (2000) (see observers #3 and #4 in Fig. 7b). As shown in Fig. 7a for direction tuning and Fig. 7c for speed tuning, the model’s predictions for many parameter combinations (points) fall within the small range empirically reported (dotted gray, horizontal lines). Under certain conditions, such as moderate sensory uncertainty (10x the fitted value), the model predicts near-zero shifts (Fig. 7b and d). Interestingly, for center speed tuning, the model can even predict shifts in the opposite direction of a simple surround-relative prediction (first and third panels in Fig. 7d, bottom row), precisely because motion is referenced to the inferred intermediate frame.

Thus, the data from Born (2000) do not rule out relative motion coding in MT. They may challenge the assumption of a fixed, pure surround-based reference frame. However, even this conclusion is limited by the small number of neurons and sparse stimulus sampling used in the experiments. Importantly, our model provides a way to resolve this ambiguity. It generates specific, quantitative predictions for the expected shift based on an observer’s behavioral parameters and their neuron’s baseline tuning. In fact, for some observers from Shivkumar et al. (2025), the minimal shifts reported by Born (2000) are exactly what our Bayesian causal inference model would predict. This highlights the need for such a principled model to guide new, densely-sampled experiments that can definitively distinguish among competing theories of motion encoding by measuring both behavioral and neural responses to the same set of stimuli.

## 3 Discussion

In this study, we demonstrated that a normative Bayesian causal inference model of motion perception can qualitatively account for the diverse center-surround interaction effects reported in the primate middle temporal area (MT). By deriving comprehensive predictions from a model fitted to human psychophysical data, we showed that the diverse, and often seemingly contradictory, response properties of MT neurons can be understood as principled manifestations of a single underlying computation: inferring the latent causal structure of the visual world. This framework unifies disparate findings in motion perception and neurophysiology, reinterprets long-standing debates about neural coding in MT, and provides a clear, theory-driven path for future experiments.

### 3.1 A single model for diverse neural effects

The function of center-surround interactions in area MT has been a puzzle, primarily due to the diversity of the observed phenomena. Our framework reframes this diversity not as a catalogue of many distinct neuron types with fixed properties, but as the consequence of a single inferential process: Bayesian causal inference.

Classic neurophysiological studies categorized the variety of surround modulations. A small subset of MT neurons (∼8%) shows no surround modulation at all, consistent with the traditional view of MT encoding motion in purely retinal coordinates (Inaba et al., 2011, 2007; Newsome et al., 1988). Our model naturally accounts for these neurons through the retinocentric latent variable (**v**_center_). The most common class of neurons exhibits direction-selective antagonistic suppression, strongest when center and surround move together (Allman et al., 1985a; Born, 2000; Born & Bradley, 2005; Born & Tootell, 1992; Tanaka et al., 1986). Our model robustly reproduces this effect as a signature of relative motion coding (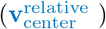) with respect to the inferred reference frame.

Crucially, the model also explains phenomena that have challenged simpler descriptive or mechanistic models. For instance, reinforcing modulation (facilitation), where responses are enhanced by same-direction surrounds, hwas been diffcult to reconcile with a purely suppressive mechanism like classic divisive normalization, in which surround activity is pooled into the normalization signal that divides the center response, so that it can only reduce, never enhance, that response (Carandini & Heeger, 2012). Our framework predicts this exact effect for neurons encoding retinocentric motion (**v**_center_) under conditions of moderate to high sensory uncertainty, providing a normative explanation for its emergence. Our model also resolves the key finding from Born (2000) that center tuning curves modulate their gain but do not shift their peak in response to surround motion – a result that poses a direct challenge to any pure surround-relative coding scheme. The causal inference model predicts this when there is larger uncertainty about the causal structures, and hence ambiguity about the correct reference frame. Because the inferred reference is often intermediate – not locked to the surround – the posterior belief about the center’s motion is modulated while its peak shifts only minimally, in line with empirical observations.

Which latent variable a neuron encodes (**v**_center_ or 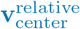) can be determined by which of the two predictions best fits its center-surround responses. If no neurons are well-explained by our model, then this could be because the neurons do not represent posteriors over the latent variables in our generative model, or because their representation is nonlinear (e.g. forming a probabilistic population code Haefner et al., 2024; Ma et al., 2006), thereby falsifying our model. If only some neurons are not well-explained by our model, then they may serve auxiliary roles in computing the posterior (Pecevski et al., 2011).

Our framework not only reinterprets past findings but also generates new, testable predictions. The model makes directly testable predictions for surround modulation. Facilitation, for instance, is predicted for neurons encoding retinocentric motion (**v**_center_) under moderate to high sensory uncertainty. In contrast, suppression is the hallmark of neurons encoding relative motion (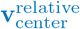) with respect to a surround-influenced reference frame. Furthermore, the model explains how direction-dependence arises from sensory noise: high uncertainty leads to broad, direction-independent modulation, whereas low uncertainty produces sharply tuned, direction-specific effects. Finally, our method directly links an individual’s perception (via behaviorally fitted model parameters) with their neural encoding (via their recorded baseline tuning profile) to produce these specific predictions. By making explicit predictions for tuning curves and firing rate variability across a broad range of center-surround stimuli, our work moves beyond explaining prior studies – which are insuffcient to rule out competing relative motion models – and provides a clear path forward for targeted new experiments.

### 3.2 Reinterpreting MT: from motion filters to causal inference

Our findings suggest a conceptual shift in our understanding of MT’s role. In traditional models of neural activity, center-surround processing in motion perception arises from a cascade of linear and nonlinear computations that integrate local motion signals into the representation of a more global motion of the visual scene (Adelson & Bergen, 1985; Albright, 1984; Barth & Watson, 2000; Movshon et al., 1985; Rust et al., 2006; Simoncelli & Heeger, 1998; Wilson et al., 1992). In these models, V1 simple cells function as linear space-time filters, detecting only local motion energy within their passband, without encoding the global motion in the scene. The role of V1 complex cells remains debated: some models suggest they compute local averages of V1 activity to represent phase-insensitive motion energy (Adelson & Bergen, 1985; Albright, 1984; Movshon et al., 1985; Rust et al., 2006; Simoncelli & Heeger, 1998), while others propose they already signal global motion direction by selectively responding to endpoints of long contours, thereby avoiding the aperture problem (Pack et al., 2003a; Pack et al., 2003b; Zarei Eskikand et al., 2016). For MT, most models posit that its cells integrate local motion energy signals from V1 and apply nonlinear computations, such as divisive normalization and half-squaring, to compute the global motion in the visual scene (Adelson & Bergen, 1985; Albright, 1984; Movshon et al., 1985; Rust et al., 2006; Simoncelli & Heeger, 1998). Alternative theories argue that nonlinear computations, such as texture boundary motion and feature extraction, occur earlier in V1 and V2, where global motion in the visual scene may already be represented. In this framework, MT primarily pools these inputs and reduces noise (Barth & Watson, 2000; Wilson et al., 1992; Zarei Eskikand et al., 2019, 2020, 2016). While these traditional models successfully explain many aspects of neural activity along the V1–MT pathways, these models are fundamentally descriptive; they explain how a computation might be implemented but not why it takes the particular form it does. They struggle to explain the functional purpose behind the diverse center-surround effects and do not clarify how the brain determines the appropriate reference frame for motion integration or segmentation. Consequently, these models fail to explain center-surround effects in perception.

In contrast, our model assumes that neural activity represents posterior beliefs about latent variables in a generative model of retinal motion. These latent variables represent not only the velocities of moving objects but also the reference frames within which the velocities are represented, both of which are inferred from sensory input. The resulting relationships between the 4-dimensional stimulus (center and surround motion, each with speed and direction) and neural responses are complex and diffcult to capture with a feedforward model. Furthermore, these relationships are dynamically modulated by contextual variables that influence the brain’s belief about causal structure, such as stimulus uncertainty (number of elements and/or contrast) and the distance between center and surround. These variables help account for reported differences between studies.

Why should the brain represent motion relative to moving reference frames? For a stationary observer (including stationary eyes), retinal motion results from objects moving in the world, and these objects are often hierarchically organized, e.g., a hand that is part of a person walking up a moving escalator. The retinal motion of the hand is then the sum of all relative velocities: the hand relative to the body, the body relative to the escalator, and the escalator relative to the world. In principle, the brain could represent every visual element in retinal coordinates, much like a flow field. However, decades of psychophysical studies have shown that this is not the case (Bill et al., 2020; Braddick, 1993; Johansson, 1950, 1973; Shivkumar et al., 2025; Shum & Wolford, 1983). Two normative principles have been proposed to explain these findings. The first is Bayesian causal inference, a special case of the Bayesian brain hypothesis (Knill & Pouget, 2004; Yuille & Kersten, 2006), which posits that the brain inverts the process that generated its observations. An observer that inverts this generative process must estimate motion relative to inferred reference frames, which is what makes motion segmentation, object grouping, and the discounting of self-motion possible. This is the classical vector-analysis view of motion perception (Johansson, 1950), recently formalized as hierarchical causal inference (Bill et al., 2022, 2020; Gershman et al., 2016; Shivkumar et al., 2025; Yang et al., 2021). The second normative principle is the minimum description length principle(Attneave, 1954; Hochberg & McAlister, 1953): because relative velocities are exactly or nearly zero most of the time, describing a scene by a shared reference frame plus small relative velocities is more compact than describing the retinal velocity of each element independently. This is precisely what the delta component of our prior expresses in the model.

Empirically, however, the hypothesis that the brain represents motion relative to moving reference frames is supported mainly by behavior. The most direct neural evidence to date is that MT responses are biased by peripheral optic flow in a way that partially accounts for perceptual flow-parsing biases, via a mechanism distinct from conventional surround suppression (Peltier et al., 2024). Our predictions provide a way to test this reference-frame hypothesis more directly.

Therefore, our results reframe MT not merely as a sophisticated filter for integrating V1 signals using simple, fixed rules, but as an inferential system that deduces the relationships between moving objects to form coherent perceptual representations. This perspective not only provides the missing functional role (“why”) for the complex repertoire of center-surround interactions – suggesting they emerge as a solution to a causal inference problem – but can also explain a wide range of empirical tuning functions.

### 3.3 Bridging perceptual phenomena and neural circuits

A central contribution of our work is bridging the gap between perceptual studies of motion and their neural underpinnings. Decades of psychophysical research have established two dominant effects in motion perception: integration and segmentation (Braddick, 1993; Dakin & Mareschal, 2000; Penaloza et al., 2024; Tadin & Lappin, 2005; Tadin et al., 2003; Zarei Eskikand et al., 2024, 2019). Integration is thought to improve the signal-to-noise ratio, especially in low-contrast conditions (Britten & Heuer, 1999; Britten et al., 1993; Tadin et al., 2003), while segmentation enhances the perception of relative motion, aiding in object tracking and parsing scenes relative to a reference frame (Huang et al., 2007; Johansson, 1950; Tadin et al., 2019). MT neurons seem to exhibit shifting responses between integration and segmentation in a dynamic, stimulus-dependent way (Huang et al., 2007, 2008).

Our results provide a unifying explanation for these observations, at both the perceptual and neural levels. The causal inference model casts both integration and segmentation as outcomes of probabilistic inference over latent motion structures (Penaloza et al., 2024; Shivkumar et al., 2025; Yang et al., 2021). We show that MT neurons can exhibit response properties consistent with either computation by representing posterior beliefs over latent variables that flexibly change with stimulus context. Importantly, perceptual states and surround modulation are not in one-to-one correspondence: the same stimulus can suppress neurons representing 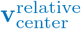 and facilitate neurons representing **v**_center_, and the same neuron can switch between suppression and facilitation depending on the stimulus (Sections 2.5.2 and 2.5.4). This offers a principled account of how the same population of neurons can support both perceptual states (integration and segmentation) and why certain center-surround stimuli elicit ambiguous or mixed neural responses.

Our model is formulated at the computational level and makes no assumptions about its neural implementation. At the circuit level, divisive normalization is often proposed as a canonical computation for center-surround effects (Carandini & Heeger, 2012; Coen-Cagli et al., 2015). While simple divisive normalization models cannot capture the full range of effects predicted by our framework, extended variants that include multiplicative interactions within their normalization pool may provide a mechanistic approximation (Penaloza et al., 2023). Several mechanistic models have been proposed to explain the integration and segmentation properties of MT neurons (Kim & Wilson, 1997; Qian et al., 1994a, 1994b; Wilson & Kim, 1994; Zarei Eskikand et al., 2019, 2020). These models generally fall into three categories: (1) those attributing segmentation to inhibitory interactions between direction-tuned neurons (Kim & Wilson, 1997; Qian et al., 1994a, 1994b; Wilson & Kim, 1994), (2) those proposing distinct populations of neurons specialized for integration versus segmentation (Beck & Neumann, 2011; Zarei Eskikand et al., 2020), and (3) those positing a single, adaptive population that dynamically adjusts its center-surround interactions based on stimulus context (Zarei Eskikand et al., 2019). While these models can account for many center-surround interactions, they struggle to explain cases that contradict simple relative motion encoding (e.g., Born, 2000; Tanaka et al., 1986). It also remains unclear whether circuit models based on divisive normalization or stabilized supralinear networks (Rubin et al., 2015) can support the probabilistic computations required by Bayesian causal inference. Building on Hennequin et al. (2014) and Echeveste et al. (2020), future work could explore this connection, using the detailed predictions from our framework as a clear computational target for mechanistic modeling.

This computational-level formulation also does not exclude auxiliary neurons whose activity is not read out downstream but is needed for the computation. For instance, in mouse V1, surround-facilitated somatostatin-expressing inhibitory interneurons (SOMs) are required for the surround suppression of pyramidal neurons (Adesnik et al., 2012), and network implementations of sampling-based inference may similarly rely on auxiliary neurons (Pecevski et al., 2011). However, our model predicts facilitation also for neurons that represent the posterior itself, whose activity is read out by downstream areas. A testable prediction is therefore that at least some surround-facilitated MT neurons project outside MT.

### 3.4 Relation to other accounts of center-surround modulation

Our claim here concerns neurons representing visual motion: for these neurons, causal inference over reference frames predicts specific patterns of center-surround interactions. We compared our model predictions to data from area MT because most primate center-surround motion experiments, and the tuning data needed to constrain the model, come from this area. Other forms of center-surround processing, e.g., in the retina or for non-motion features, would require their own generative models.

The closest example of such a model is the normative account of surround modulation in V1 (Coen-Cagli et al., 2012, 2015). Their account and ours share the core underlying logic: contextual modulation is a consequence of inference about scene structure, and the surround influences the center only to the extent that the two are inferred to belong together. In both cases, the model is also constrained independently of the neural data, by natural image statistics in their case and by human psychophysics in ours. The accounts differ, however, in their latent variables. In the V1 model, inference determines whether center and surround filter responses share a common gain (normalization) signal. In our model, the latent variables are velocities, the physical quantities that the observer seeks to estimate. Our model therefore infers not only whether center and surround are related, but also the reference frame relative to which the center’s motion is represented, a latent variable with no analogue in the V1 model. This additional variable allows facilitation to arise in structured regions of stimulus space, whereas facilitation is rarer in the V1 model. We therefore see the two accounts as complementary: the same general principle, applied to different features in different areas.

Interestingly, V1 neurons responding to drifting gratings show a form of surround suppression similar to the one we predict for neurons representing relative motion, 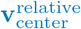, in our model: direction-selective suppression that is strongest when the surround moves in the neuron’s preferred direction and decreases gradually as the surround direction deviates from it (Cavanaugh et al., 2002; Walker et al., 1999). We mention this resemblance only as an observation, not as a test of the model. A proper test in V1 would require a generative model adapted to these stimuli.

Center-surround interactions also emerge in deep feedforward networks trained on object recognition, without any explicit causal inference objective (Pan et al., 2023). We see no conflict between this finding and our account. In natural images, context is informative about the identity of objects, so a normative account predicts that networks optimized for object categorization should develop contextual modulation (Oliva & Torralba, 2007).

### 3.5 Limitations and Future Directions

This study provides a qualitative, proof-of-principle demonstration that a normative model of perception can account for a wide range of neurophysiological data. Three main limitations, however, point toward crucial avenues for future research.

First, linking Bayesian models to neural activity requires an assumption about the neural code. We used the neural sampling hypothesis (e.g., Buesing et al., 2011; Fiser et al., 2010; Haefner et al., 2016; Hoyer and Hyvärinen, 2002; Orbán et al., 2016), which belongs to a broad class of Linear Distributional Codes (LDCs) that also includes Distributed Distributional Codes (DDCs) (Lange & Haefner, 2022). While our predictions for mean tuning curves are generalizable across LDCs (see Methods 4.5 and Lengyel et al., 2023), our predictions for response variability are specific to sampling-based codes. The broader debate over whether the brain uses LDCs or other encoding schemes, like Probabilistic Population Codes (PPCs), is ongoing (Beck et al.,2012, 2020; Fiser et al., 2010; Haefner et al., 2024; Lange et al., 2022; Ma et al., 2006; Pouget et al., Tajima et al., 2016; Ujfalussy & Orbán, 2022; Vértes & Sahani, 2018, 2019). A critical next step is to derive predictions for non-LDC codes like PPCs, which would generate directly competing and testable hypotheses.

Second, our comparison to historical data is qualitative. While we show that observed patterns of neural activity are broadly consistent with our model, rigorous quantitative model comparison against alternatives (like extended divisive normalization models) is required. Moreover, the neural data we compared against were recorded in anesthetized monkeys (Allman et al., 1985a; Born, 2000; Tanaka et al., 1986), whereas our model parameters were fitted to the behavior of awake human observers. Because our predictions depend on observer-specific parameters, such as sensory noise and the prior probability of grouping, which may change with anesthesia, this comparison is necessarily imperfect. Although surround suppression has also been observed in alert monkeys (Hunter & Born, 2011), the narrower stimulus sampling does not allow testing the full range of predicted effects. The data we used are therefore the most suitable currently available, but are far from ideal for testing the model.

Importantly, our library of neural predictions contains no parameters fitted to neural data: it follows directly from model fits to human psychophysics (Shivkumar et al., 2025) and from center velocity-only (with stationary surround) tuning curves measured from area MT in passively fixating monkeys (DeAngelis & Uka, 2003). We then only selected the best-matching prediction from this fixed library for each empirical class of center-surround interaction. This transfer from human perception to MT physiology captures diverse center-surround effects, which is unlikely to be coincidental and suggests a shared computational principle. To our knowledge, Bayesian causal inference over reference frames is currently the only normative account that unifies these neurophysiological findings. A decisive quantitative test would require fitting the model parameters to an animal’s behavior while simultaneously recording from MT neurons, and then predicting each neuron’s center-surround responses from its own center-only (i.e., with stationary surround) tuning data. Our framework is ideally suited to guide such work, as it can generate predictions for any stimulus, allowing researchers to design optimized experiments that can maximally differentiate between competing models (i.e., controversial stimuli; Golan et al., 2020).

Third, as for any Bayesian model, our predictions depend on whether the prior and the likelihood are well-specified. The form of the prior is motivated by the physics of the world rather than fitted freely in shape. Its delta component reflects that, due to friction, objects are mostly stationary relative to their reference frame (Shivkumar et al., 2025), and its Gaussian component is the slow-speed prior that explains many motion illusions (Weiss et al., 2002). This form of the prior is not the only possible choice: when recovered from human speed judgments, the slow-speed prior has heavier tails than the Rayleigh distribution over speed implied by our two-dimensional Gaussian prior over velocity (Stocker & Simoncelli, 2006), and combining the delta component with a heavier-tailed distribution over velocity would better capture perception at high speeds (Shivkumar et al., 2025). The likelihood is assumed to be an isotropic Gaussian. Only these functional forms are shared across observers; their parameters, such as the width of the slow-speed prior and the sensory noise, were fitted separately to each observer’s behavior (Shivkumar et al., 2025).

Note that our predictions concern posteriors, so they do not require MT to represent the prior explicitly: the prior could be implemented upstream, through feedback, or implicitly in the connectivity of recurrent circuits (Echeveste et al., 2020).

The parameter estimates of Shivkumar et al. (2025) also differ in how tightly they are constrained by behavior (Supplementary Fig. S5 in Shivkumar et al., 2025). However, the main signatures of relative motion coding, direction-selective surround suppression and the diagonal interaction, were robust across all five observers (Fig. S2), across a hundredfold range of sensory uncertainty (Fig. 6 and Fig. S5), and across a broad sample of parameters around the fitted values (Fig. S4).

Finally, our prior is itself a hypothesis about the statistics of natural scenes: due to friction elements are usually exactly stationary relative to their natural reference frame and otherwise tend to move slowly. This links our framework to accounts that explain center-surround effects in V1 by adaptation to natural image statistics (Coen-Cagli et al., 2012; Rao & Ballard, 1999; Schwartz & Simoncelli, 2001). Our prior is defined over latent velocities in reference frames, whereas measured statistics describe the observed retinal image motion. Therefore, we would need to integrate out the latent variables of our model (possibly together with other important latent variables that we would need to add to the model such as object distance) to get a distribution over retinal image velocity that could be compared directly with natural videos. We expect this to resemble the scale-mixture structure recently reported for natural motion (Salisbury & Palmer, 2025). A mismatch between the prior that best explains perception or neural responses and these statistics would then point to a misspecification of the model, and how to improve it.

### 3.6 Conclusion

This study establishes Bayesian causal inference as a unifying normative principle for center-surround interactions in motion. Our framework parsimoniously explains a wide array of disparate findings – from surround suppression and facilitation to minimal tuning curve shifts in response to surround motion – as the principled outputs of a single computation over latent reference frames. By reinterpreting classic neurophysiological data and bridging the long-standing gap between models of perception and neural activity, this work reframes our understanding of MT’s role from that of a motion filter to that of an inference engine. This provides a clear foundation for designing new, theory-driven experiments to dissect the circuit-level implementation of these abstract cognitive computations.

## 4 Methods

### 4.1 The behavioral task and data source

The Bayesian causal inference model was constrained using behavioral data from five observers in Experiment 1 of Shivkumar et al. (2025). In that study, observers used a dial to report their perceived motion direction of a central patch of green dots surrounded by a peripheral patch of red dots (Fig. 2a).

The surrounding dots were either stationary or moved horizontally (0 or 180°) at a speed (*v*_0_) of 1 deg./sec. Stimuli were displayed at an eccentricity of 5° in the periphery. On each trial, the direction of the central patch, Δ*θ*, was randomly selected from the set 0°, ±2.5°, ±5°, ±10°, ±20°, ±45°. The center always moved with the same horizontal velocity component, *v*_0_, as the moving surround (Eqs. 5 and 6), so that with a moving surround, the center’s velocity relative to the surround was always vertical, i.e., *±*90^*°*^ depending on the sign of Δ*θ*. With a stationary surround, the center’s relative velocity equals its retinal velocity. After a fixation period of 0.5 seconds, the stimulus appeared and its velocity reversed every 1.5 seconds, following a square-wave profile with a period of 3 seconds, which kept the patches within a fixed area of the screen. The stimulus timing is described by a square wave, *q*(*t*), where *t* = 0 denotes trial onset:

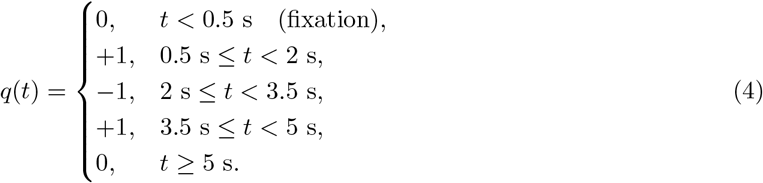

With **e**(*ψ*) = (cos *ψ*, sin *ψ*)^*⊤*^ denoting a unit vector in direction *ψ*, the surround velocity is

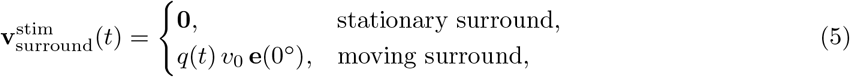

and the center velocity is

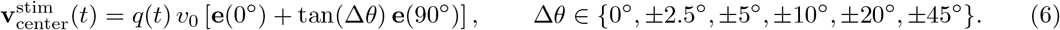

Thus, the center moves in direction Δ*θ* with speed *v*_0_*/* cos Δ*θ*, and its velocity relative to the moving surround, *q*(*t*) *v*_0_ tan(Δ*θ*) **e**(90^*°*^), is always vertical. For a detailed description of the stimuli (including videos), task, and design, see Shivkumar et al. (2025).

When the surround was stationary, reported directions closely followed the true motion of the center patch as displayed on the screen. However, when the surround patch was moving, observers’ responses deviated systematically from the true motion of the center patch. For small center directions, responses were biased towards zero degrees (the surround direction), consistent with observers integrating the center and surround velocities. For larger center directions, responses were biased towards 90°, indicating that observers perceived the relative velocity between the center and surround. Additional findings from this experiment, along with further analyses, are detailed in Shivkumar et al. (2025).

### 4.2 The Bayesian causal inference model

The Bayesian causal inference model described in Shivkumar et al. (2025) was used to derive posterior beliefs for center-surround motion stimuli. Here, we summarize its generative model while the derivation of the inference equations and the model fitting procedure are given in Shivkumar et al. (2025).

#### Generative model

The retinal velocity of each visual element (i.e., the center and surround patches) is the sum of the reference frame’s velocity to which the element belongs and its relative velocity with respect to that reference frame.

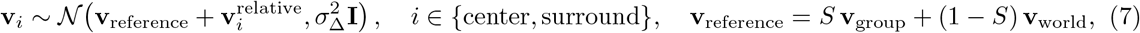

where 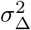 is the variance representing computational noise, and *S ∈ {*0, 1*}* indicates whether the center and surround share a common moving reference frame (referred to here as the group, **v**_group_), with prior probability *p*(*S* = 1) = *β*. Otherwise, both patches move in the stationary world frame (**v**_world_ = **0** for a stationary observer). The group velocity (i.e., the common moving reference frame) is itself defined relative to the world, 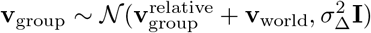).

The relative velocity of each element is either exactly zero (stationary within its reference frame) or slow. We formalize this with a binary variable, *C*_*j*_, which indicates whether element *j ∈ {*center, surround, group} moves within its reference frame:

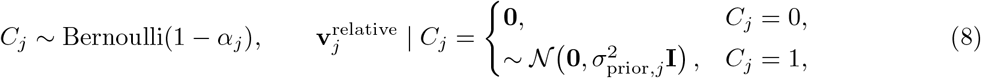

Marginalizing over *C*_*j*_ yields a mixture prior over each relative velocity,

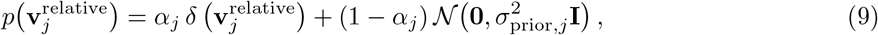

where the delta component represents elements that are stationary relative to their reference frame, and the Gaussian component represents slow motion within it. Consequently, this prior naturally enables the model to infer whether elements are stationary relative to their reference frame or moving within their reference frame.

#### Observation model

The observed retinal velocities are assumed to be the true retinal velocities corrupted by sensory noise (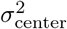 and 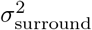)

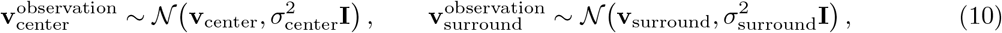

where all velocities are two-dimensional vectors and **I** is the 2 *×* 2 identity matrix.

#### Causal structures and inference

A causal structure, *s* = (*S, C*_center_, *C*_surround_, *C*_group_), specifies whether the patches are grouped (*S*) and, for each relative velocity, whether it is drawn from the delta (*C*_*j*_ = 0) or the Gaussian (*C*_*j*_ = 1) component. This gives 2^3^ = 8 structures with grouping and 2^2^ = 4 without, i.e., 12 in total. Given a structure, the model is linear-Gaussian, so the posterior over each latent velocity is computed in closed form and then combined across structures (Equation (2)), with *p*(*s* |**v**^observation^) ∝*p*(**v**^observation^ |*s*) *p*(*s*).

#### Perceptual read-out

The perceived center velocity is the non-zero relative velocity lowest in the hierarchy (Shivkumar et al., 2025):

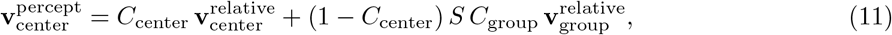

and the reported direction is the direction of this vector. The model parameters were fitted to these reports (Shivkumar et al., 2025).

We adopted a simplified version of the Bayesian causal inference model in Shivkumar et al. (2025) which includes 10 parameters, fitted separately for each observer: (1) two sensory uncertainty parameters associated with the center and surround patches’ observed velocities (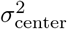 and 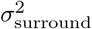), (2) one computational noise parameter 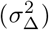, (3) three mixture prior parameters corresponding to the group, center, and surround velocities (*α*_*j*_), (4) three parameters for the prior widths of the zero-mean Gaussian slow speed priors over the group, center, and surround velocities (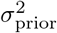), and (5) one parameter representing the probability of grouping the center and the surround patches into a single causal structure (*β*). Shivkumar et al. (2025) obtained full posterior distributions over these parameters (their Supplementary Fig. S5); here we used the fitted values for each observer, and we explored a broader range of values around them in Methods 4.6 (Fig. S4). Detailed descriptions of these parameters and their fitted values can be found in Shivkumar et al. (2025).

#### Constraint and identifiability

The prior and likelihood parameters were fitted jointly to each observer’s direction reports (Shivkumar et al., 2025). The parameters cannot be separated from a single stimulus condition, but the experimental design of Shivkumar et al. (2025) constrains them separately: it includes a stationary-surround control and a range of center directions, across which the sensory noise mainly determines the spread of the responses, whereas the prior parameters mainly determine the shape of the bias as a function of the center-surround difference. Remaining trade-offs between parameters would appear as correlations in their joint posterior, which Shivkumar et al. (2025) report in their Supplementary Fig. S5.

The Gaussian form of the likelihood with a constant variance is a simplification: human speed perception suggests that sensory noise grows with speed (Stocker & Simoncelli, 2006) (following Weber’s law), but this has little effect over the narrow range of speeds used in the experiment.

### 4.3 Model fitting results from Shivkumar et al. (2025)

The fitted mixture weights (*α*_*j*_) placed most of the prior mass on the delta component, reflecting the brain’s strong expectation that relative motion in the world is exactly zero, rather than merely slow. Moreover, observers attributed significant probability mass to only four out of the 12 possible causal motion structures:

1. Motion integration: Integrating center and surround velocities, leading to the perception of cue-combined motion velocity (Fig. 2c). The posterior probability of this structure was highest when the center and surround moved at the same velocity and decreased as the separation between center and surround velocities increased (Fig. S1c and d).
2. Surround reference: Perceiving center motion in the reference frame defined by the surround (i.e., pure relative motion or motion segmentation, see Fig. 2d). This structure was prominent for intermediate differences between center and surround motion directions for a subset of observers (Fig. S1e and f).
3. Center reference: Perceiving surround motion in the reference frame of the center. In this case, the perception of center motion largely aligned with its retinal motion (Fig. 2e), with only a few observers attributing significant probability mass to this structure (Fig. S1g and h).
4. Intermediate reference: Perceiving both center and surround motion relative to a reference frame intermediate between the two (Fig. 2f). This structure accounted for incomplete subtraction of the surround velocity from the center velocity at larger differences between center and surround motion directions and was prominent among most observers (Fig. S1i and j).

Notably, over the range of stimuli tested by Shivkumar et al. (2025), none of the observers attributed substantial probability mass to the possibility that the center and surround patches were independent, indicating that they did not perceive the two patches as causally unrelated (Fig. 2g and Fig. S1 k and l).

Interestingly, motion structures with intermediate reference frames have higher posterior probabilities than other structures at large separations. This is due to the Gaussian slow-speed component in the prior, which favors smaller relative velocities under an intermediate reference frame, outweighing the mass concentrated in the delta component of the prior. Additional insights from fitting the model to observers’ responses, along with further analyses, can be found in Shivkumar et al. (2025).

### 4.4 Generating neural predictions from Bayesian models

We generated neural predictions from the model’s posterior beliefs, assuming that they are represented in the single-neuron activity via Neural Sampling (Fiser et al., 2010; Hoyer & Hyvärinen, 2002).

We used the following algorithm to link a posterior distribution to mean firing rates and variability in the firing rates:

1. Draw *N* samples from the posterior over the latent variable *k* represented by the neuron, given the center-surround stimulus 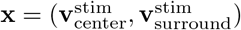:

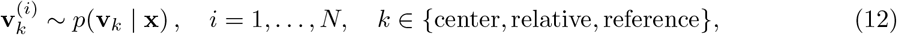

where **v**_center_, 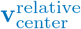, and **v**_reference_ are defined in Methods 4.2, and each sample 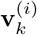 has speed *u*^(*i*)^ and direction *θ*^(*i*)^.
2. Map each sample to an instantaneous firing rate using the neuron’s tuning curve measured with a stationary surround:

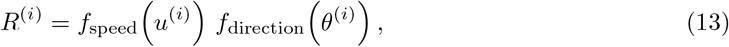

where *f*_speed_ and *f*_direction_ are the neuron’s speed and direction tuning functions.
3. Compute the predicted mean firing rate and its variability:

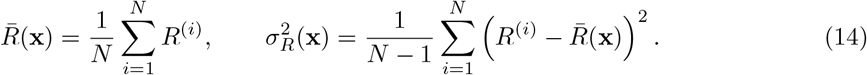
4. Repeat for each center-surround stimulus **x**.

All predicted tuning curves (mean firing rate) and response variability (variance of the firing rates) presented in this study were generated using the algorithm described above. This variability term quantifies the uncertainty in the model’s posterior beliefs and represents the predicted “excess variance” (over-dispersion) that modulates a baseline Poisson process, accounting for super-Poisson spike count statistics (Goris et al., 2014).

### 4.5 Predicted tuning curves hold for all Linear Distributional Codes

The method described above can be regarded as a special case of the approach introduced in Lengyel et al. (2023). In that work, Lengyel et al. (2023) developed a method for testing Bayesian models using neural activity, which applies to all Linear Distributional Code (LDC; Lange and Haefner, 2022) based encoding models. This approach also enables researchers to generate predicted tuning curves for single neurons that are consistent across all LDCs (Lange and Haefner, 2022). LDCs encompass a broad class of encoding models, including schemes within two of the three principal families of hypotheses about neural encoding: Neural Sampling Codes (Fiser et al., 2010; Hoyer & Hyvärinen, 2002) and Distributed Distributional Codes (DDCs; Vértes and Sahani, 2019). Consequently, the neural predictions derived from this method hold for any Neural Sampling or DDC-based encoding used by the brain to represent probability distributions. However, the method does not apply to Probabilistic Population Codes (PPCs; Ma et al., 2006) or to other nonlinear distributional codes such as expectile codes (Dabney et al., 2020); see Lengyel et al. (2023) and Lange and Haefner (2022) for details.

In essence, the approach of Lengyel et al. (2023) identifies “test” stimuli whose posteriors can be expressed as mixtures of the posteriors elicited by other, “baseline” stimuli. LDCs are defined by the property that the mean response of a neuron to a mixture of distributions equals the same mixture of its mean responses to the individual distributions (Lange & Haefner, 2022). Consequently, if the posterior for a test stimulus is a mixture of the posteriors for a set of baseline stimuli, the neuron’s mean response to the test stimulus is the same mixture of its mean responses to the baseline stimuli, with the same mixture weights (see the derivation in Lengyel et al., 2023).

The method we used to generate tuning curves (i.e., mean firing rates) is a special case of this approach, obtained under the following assumptions. The *test stimulus* is a center-surround stimulus with a moving surround, 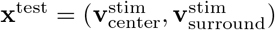, whose posterior over the latent variable represented by the neuron, *p*(**v**_*k*_ | **x**^test^), is computed with the fitted model parameters (step 1 above). The *baseline stimuli* are center-surround stimuli with a stationary surround, 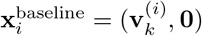, whose center velocities are given by the posterior samples 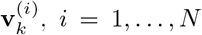. We assume that the posterior elicited by each baseline stimulus is concentrated at its center velocity, i.e., that it is well approximated by a delta function,

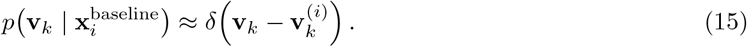

This holds when the baseline posterior is much narrower than the neuron’s tuning curve. Under this assumption, the mean response to each baseline stimulus is given by the neuron’s tuning curve measured with a stationary surround (step 2 above),

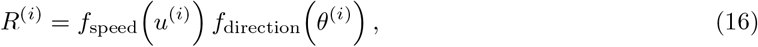

where *u*^(*i*)^ and *θ*^(*i*)^ are the speed and direction of 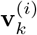. Because the samples are drawn from the test posterior, the test posterior is approximated by an equally weighted mixture of the baseline posteriors,

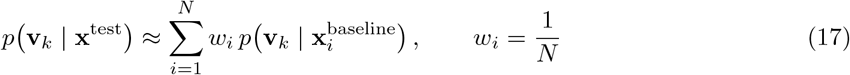

By the defining property of LDCs, the predicted mean response to the test stimulus is the same mixture of the baseline responses,

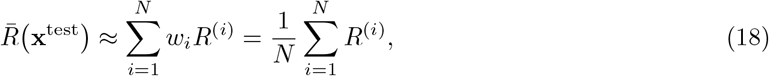

which is exactly the mean firing rate computed in step 3 (Equation (14)). As *N → ∞*, this sum converges to the expected firing rate under the test posterior, ∫*p*(**v**_*k*_ | **x**^test^) *f*_speed_(*u*) *f*_direction_(***θ***) d**v**_*k*_, where *u* and *θ* are the speed and direction of **v**_*k*_.

Therefore, the predicted tuning curves hold for any LDC, whether sampling-based or DDC-based. They do not hold for Probabilistic Population Codes (PPCs; Ma et al., 2006), which encode the natural parameters of the posterior, or for other nonlinear distributional codes, such as expectile codes (Dabney et al., 2020; Lange & Haefner, 2022). The predicted response variability, in contrast, applies only to sampling-based codes, because the LDC property constrains only mean responses, whereas under sampling, single-trial responses reflect individual posterior samples.

### 4.6 Analyzing the predicted neural activity

The center-surround stimulus described in Methods 4.1 has four important properties that modulate posterior beliefs in the Bayesian causal inference model: (1) the direction of the center, (2) the direction of the surround, (3) the speed of the center and (4) the speed of the surround patches. Therefore, we plotted the predicted tuning curves and variability as a function of these four stimulus dimensions (see Fig. 5, and Figs. S2 and S3). We then demonstrate that these predictions are different for different observers via the model parameter values fitted to their behavior (see Figs. S2 and S3). Finally, we demonstrate that these predictions are different for different neurons via their different tuning curves (see Figs. 5 and S5).

In order to investigate the variety of possible neural center-surround interactions predicted by Bayesian causal inference, we generated neural predictions using a wide range of possible model parameters besides the fitted parameters for the 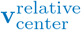 latent variable. In short, we used the five sets of parameters fitted to the five observers in Shivkumar et al. (2025) and sampled the model parameter values around the fitted values to generate 210 different predictions for each of the 9 neuron tuning profiles and for each of the 5×5 center-surround speeds and 60×60 center-surround directions. This resulted in 47250 different prediction maps across center-surround directions (210 (model parameters) x 9 (tuning profiles) x 5 (center speed) x 5 (surround speed) = 47250). The tuning profiles included three levels of widths for the speed and direction tunings from sharp to wide tuning curves (i.e., 3 speed tunings x 3 direction tunings = 9 tuning profiles, see Fig. S5). The center of the tuning curves always represented the preferred direction or speed of the neuron. Fig. S4 shows the types of center-surround interactions predicted by the causal inference model assuming 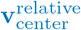 is encoded.

The main signature of our causal inference model encoding relative motion is the diagonal interaction between center and surround directions. The prediction maps in Fig. S4 are ordered from strongest (upper left corner) to weakest (lower right corner) diagonal interaction. Formally, a neuron representing motion purely relative to the surround responds as *r*(**v**_center_, **v**_surround_) = *f* (**v**_center_ − **v**_surround_), where *f* is its center-only (i.e., with stationary surround) tuning curve, whereas a surround-insensitive neuron responds as *r* = *f* (**v**_center_). Within a prediction map (fixed speeds), the latter produces vertical bands that are mirror-symmetric about the horizontal midline (surround direction = 0^*°*^), while the former is organized along the diagonal, where the velocity difference is smallest, and breaks this symmetry. Unlike a surround-insensitive response, or one in which the surround only scales the response as a function of surround direction, this response cannot be written as a product of separate center and surround tunings. We therefore measured the strength of the diagonal interaction as how mirror-symmetric the prediction map was about the horizontal midline. The lower the symmetry score, the stronger the diagonal interaction (see the scores on top of each prediction map). In the full model, however, the reference frame is inferred and the posterior is spread over several causal structures, so the predicted maps lie between the pure surround-relative case (strong diagonal) and the surround-insensitive case (vertical bands): the more observers infer the surround patch as the reference frame, and thus perceive the center motion relative to the surround, the stronger the predicted diagonal interaction. Importantly, while this library of 47250 maps was generated to explore the model’s full predictive range (Fig. S4), it was not used for the subsequent comparison with empirical data. For that validation, we only used the parameters fitted to the five observers in Shivkumar et al. (2025). This allowed us to test the specific hypothesis that the same parameters explaining human behavior could also predict the neural responses.

### 4.7 Qualitative comparison to empirical data

To qualitatively compare our model’s predictions to previously published neural data, we followed a three-step procedure. First, we generated a catalogue of potential neural responses. Second, we defined the specific stimulus conditions (“cross-sections”) needed to match each historical study. Third, for each study, we selected the best-fitting prediction from our catalogue.

#### Generation of the neural prediction catalogue

Our predictions needed to account for variability across both observers and neurons. We therefore generated a catalogue spanning two dimensions: (1) Observer Parameters: We used the five unique parameter sets fitted to the human observers in Shivkumar et al. (2025). (2) Neuron Tuning Profiles: We created nine plausible tuning profiles by combining three distinct tuning widths (approximating the 25th, 50th, and 75th percentiles from DeAngelis and Uka (2003)) for both speed and direction (3 speed widths × 3 direction widths; see Fig. S5a).

For each of the 45 combinations (5 observers × 9 tuning profiles), we generated predicted mean firing rates (tuning curves) and response variability across a dense, four-dimensional grid of center and surround speeds and directions.

#### Stimulus cross-section selection and matching procedure

To compare our predictions with the modulation effects reported in the literature, we first calculated a surround modulation index (facilitation or suppression) by subtracting the predicted response to a stationary surround from the predicted response to a moving surround (Fig. 6a).

We then extracted specific “cross-sections” from our 4D predictions to precisely match the stimulus conditions of each empirical study. For each dataset, we identified the best-fitting prediction from our 45-member catalogue by selecting the observer/tuning-profile combination that yielded the lowest root mean squared error (RMSE). The specific cross-sections were defined as follows:

1. To match Allman et al. (1985a): We fixed the center motion to the neuron’s preferred direction and speed, and varied the surround direction (0°, 30°, 60°, 90°, 120°, 150°, 180°).
2. To match Born (2000): We used a similar cross-section, varying the surround direction from 0° to 180° in 45° increments.
3. To match Tanaka et al. (1986): For direction modulation, we varied the surround direction (0°, 30°, 60°, 90°). For speed modulation, we fixed both center and surround to the preferred direction and varied the surround speed (0.25x, 1x, and 4x the preferred speed).

Other stimulus parameters, such as the type of center stimulus, stimulus size, eccentricity, dot density, and absolute speeds, also differed across the three studies and are not reported comparably. We therefore expressed all cross-sections relative to each neuron’s preferred direction and speed.

## 5 Code availability

The code that implements our analysis in the current study is publicly available in the CI neural predictions GitHub repository, https://github.com/GaborLengyel/CI_neural_predictions.

## 6 Acknowledgements

This work was supported by the BRAIN Initiative grant from the National Institute of Neurological Disorders and Stroke (https://www.ninds.nih.gov/U19NS118246 to GCD and to RMH), and by the Computing Module of a National Eye Institute Core grant (https://www.nei.nih.gov/ EY001319 to GCD). The funders had no role in study design, data collection and analysis, decision to publish, or preparation of the manuscript.

## Footnotes

1 In an experiment where participants’ heads and eyes are fixed, retinal, ego-centered, and world coordinates are indistinguishable – and our predictions do not depend on that distinction.

