## Supplementary figures for "Bayesian causal inference unifies perceptual and neuronal processing of center-surround motion in area MT"

October 24, 2026

**Abstract**

Center–surround interactions are a hallmark of visual processing and are especially prominent in area MT, where surround motion can either suppress or facilitate neuronal responses depending on context. However, existing mechanistic descriptions, including divisive normalization, do not explain the full diversity of these effects or their relationship to motion perception. Here, we show that both perceptual and neuronal center–surround phenomena can be understood as consequences of Bayesian causal inference over reference frames. Building on a normative model of motion perception, we derived predictions for the mean responses and variability of single MT neurons across the full four-dimensional space of center and surround directions and speeds. The model generates structured patterns of suppression, facilitation, and coordinate-frame selectivity that qualitatively match the diversity of center–surround effects reported in primate MT. Our results provide a unified computational account linking motion integration and segmentation in perception with contextual response modulation in MT, and yield testable predictions for how the visual system infers and represents reference frames.

**Keywords:** Bayesian causal inference; Center-surround processing; Motion perception; Neural coding in MT; Motion integration; Motion segmentation

[1] Department of Brain and Cognitive Science, University of Rochester, NY, US
[1] Center for Visual Sciences, University of Rochester, NY, US
[2] Zuckerman Institute, Columbia University, New York, NY, US
[3] Institute of Science and Technology Austria, Klosterneuburg, Austria
[†] Equal contribution

### Supplementary Material

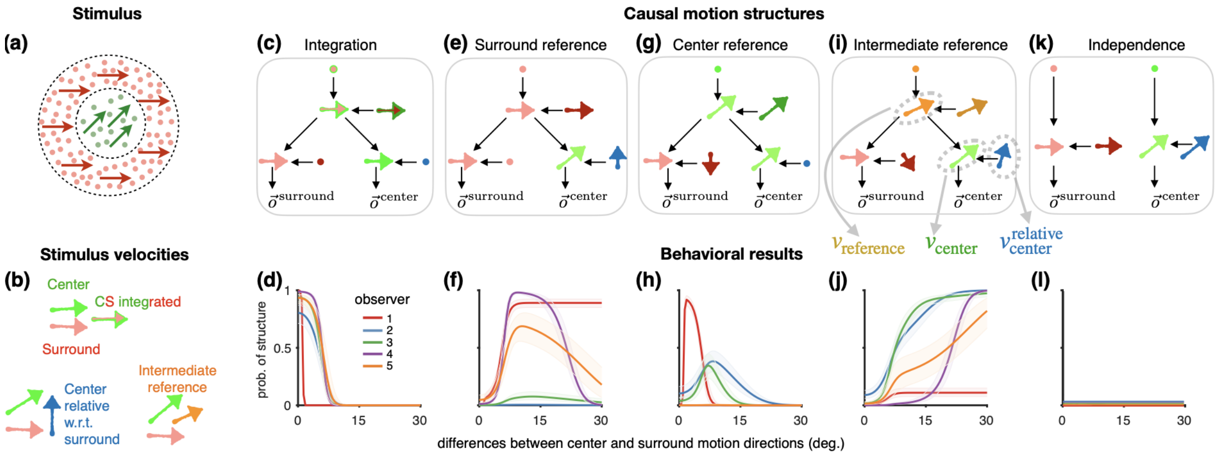


Figure S1: **Hierarchical Bayesian causal inference model of motion perception.** **(a):** Typical paradigm investigating center-surround processing in motion perception. Two patches consisting of coherently moving dots (green and red patches) are presented on the screen, and observers have to estimate the motion direction of the center patch (green). Green and red arrows represent the velocities of the center and the surround patches, respectively. **(b):** Example stimulus velocities (light red and green arrows) and the implied integrated (red-green arrow), relative (dark blue arrow), and intermediate (orange arrow) velocities. Arrow colors are matched to the colors of the variables in the generative models. **(c)–(j):** The four causal motion structures for which participants inferred significant posterior probability mass: integration (c, d), surround reference (e, f), center reference (g, h), and intermediate reference (i, j). All four assume that the center and surround patches are grouped. Top row (c, e, g, i): generative models of each structure; arrows represent non-zero velocities and points represent zero velocities (i.e., stationary). Bottom row (d, f, h, j): posterior probability (median and 95% CI) assigned to each structure by each observer (colored curves) as a function of the difference between the center and surround directions (x-axis). All these structures assume that the center and surround patches are grouped. **(k) & (l):** The causal motion structure in which the center and surround patches are independent (k), and its posterior probability, which was negligible for the stimulus range used in the experiment (l). The three latent variables contributing to the representation of the center patch's motion, $\mathbf{v}_{\text{reference}}$ , $\mathbf{v}_{\text{center}}$ , and $\mathbf{v}_{\text{center}}^{\text{relative}}$ are highlighted with gray arrows in panel **(i)**. The figures are adapted with permission from Shivkumar et al. (2025).

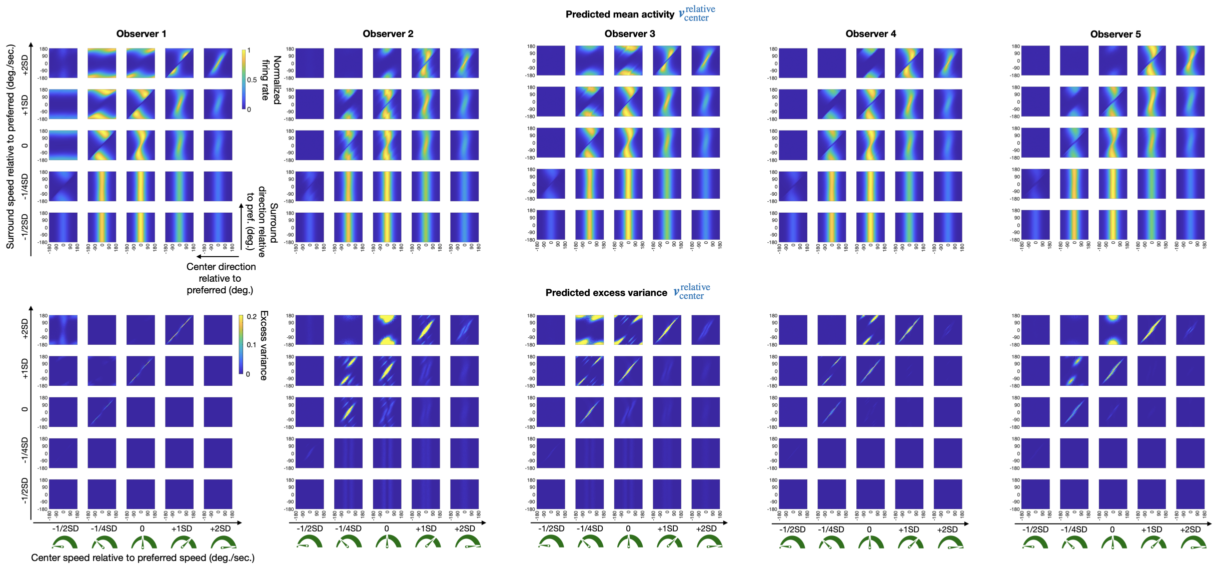


Figure S2: **Predicted 4D tuning curves for all five observers using the tuning curve of "Neuron type 5" shown in Fig. S5a**, assuming that the neuron represents the relative motion variable, $\mathbf{v}_{\text{center}}^{\text{relative}}$ .
Same as Fig. 5 but for all observers from Shivkumar et al. (2025).

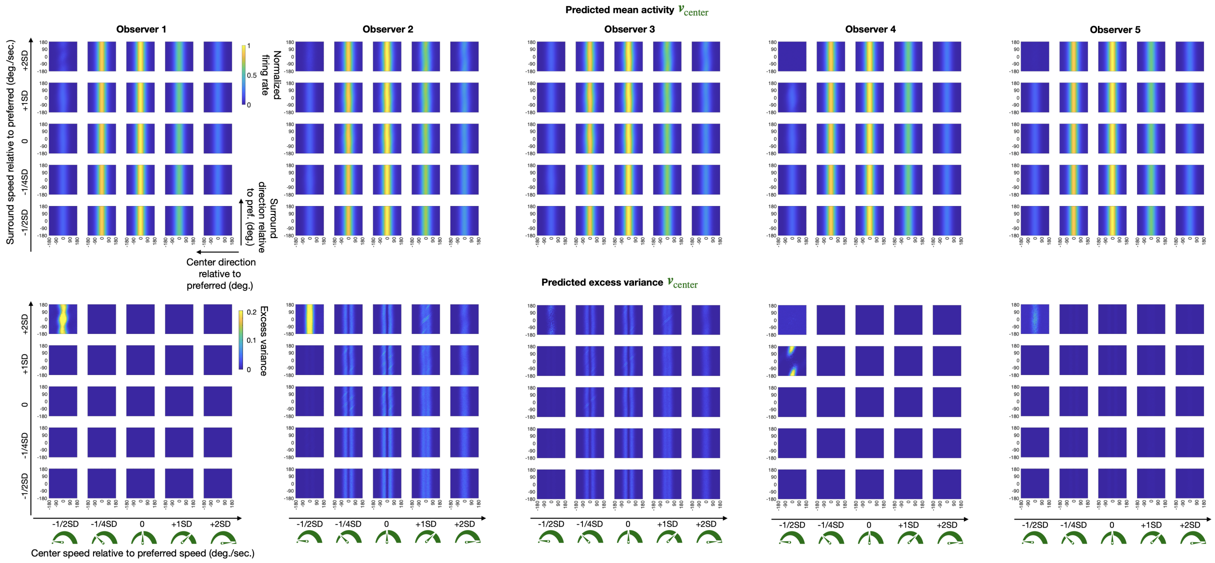


Figure S3: **Predicted 4D tuning curves for all five observers using the tuning curve of"Neuron type 5" shown in Fig. S5a**, assuming that the neuron represents the retinocentric motion variable, $\mathbf{v}_{\text{center}}$ .

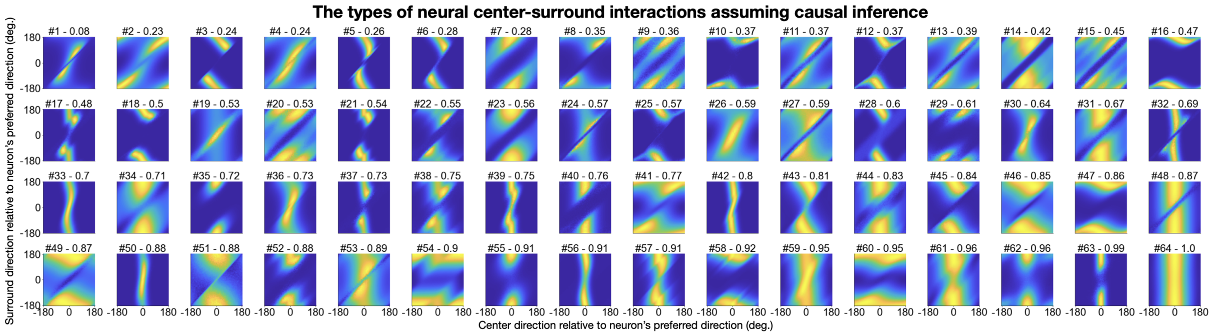


Figure S4: **Types of center-surround interactions in the mean firing rate predicted by causal inference assuming $\mathbf{v}^{\text{relative}}_{\text{center}}$ is encoded by an MT neuron with typical motion tuning (the median of empirically measured MT tuning curves from monkeys (DeAngelis & Uka, 2003)).** Shown are 64 example maps selected from the 47,250 predictions described in Methods 4.6 to be maximally different from each other, i.e., with the lowest pairwise correlations.. The predicted tuning curves are ordered from #1 - #64 by the strength of their diagonal interaction. The diagonal interaction scores are printed on top of each tuning curve panel, where 1 represents a map that is fully mirror-symmetric about the horizontal midline (no diagonal interaction) and 0 represents the strongest diagonal interaction. In short, a diagonal interaction means that the response is organized along the diagonal of each panel, where center and surround move in the same direction and the velocity difference between them is smallest. A surround-insensitive neuron would instead produce vertical bands that are mirror-symmetric about the horizontal midline (surround direction = 0°); the score quantifies the loss of this symmetry (see Methods 4.6).

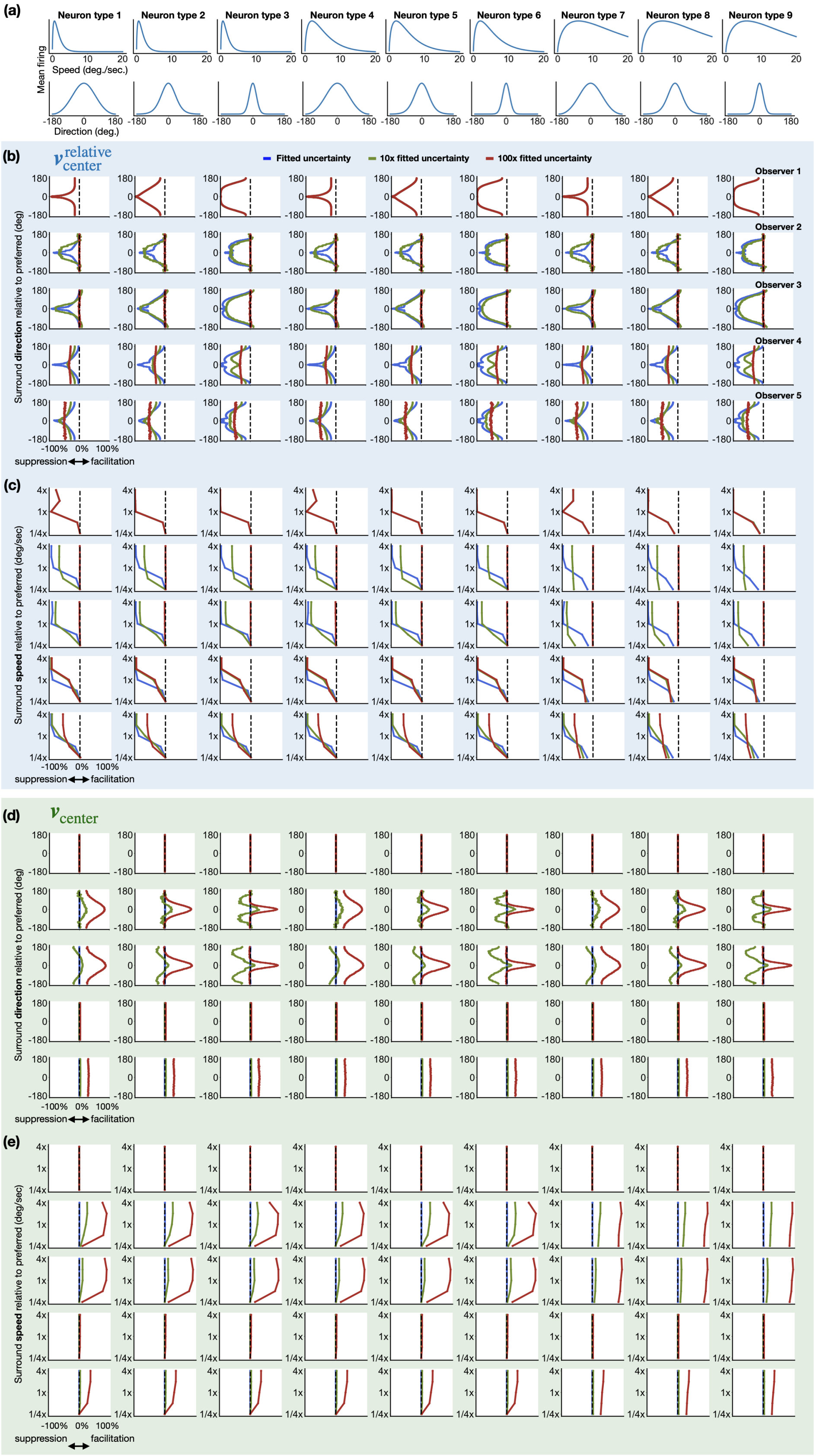


Figure S5: **Comprehensive neural predictions across observers, neuron types, and levels of sensory uncertainty.** Each panel shows predicted surround modulation (suppression/facilitation, x-axis) for all five observers (rows) and nine plausible tuning curve shapes (columns) for the range of center-surround stimuli used in previous empirical studies. Colored curves represent three levels of sensory uncertainty in the Bayesian model: the empirically fitted value (from Shivkumar et al. (2025)), 10x that value, and 100x that value. **(a)** The nine tuning curve shapes used for the predictions in the columns below. **(b)-(c)** Predictions for a neuron encoding relative motion ($\mathbf{v}_{\text{center}}^{\text{relative}}$ ). Modulation is shown as a function of surround direction in (b) and surround speed in (c). **(d)-(e)** Predictions for a neuron encoding retinocentric motion ($\mathbf{v}_{\text{center}}$ ). Modulation is shown as a function of surround direction in (d) and surround speed in (e).
